# Direct and indirect pathway neurons in ventrolateral striatum differentially regulate licking movement and nigral responses

**DOI:** 10.1101/2021.04.11.439386

**Authors:** Zhaorong Chen, Zhi-Yu Zhang, Wen Zhang, Taorong Xie, Yaping Li, Xiao-Hong Xu, Haishan Yao

## Abstract

Drinking behavior in rodents is characterized by stereotyped, rhythmic licking movement, which is regulated by the basal ganglia. It is unclear how direct and indirect pathways control the lick bout and individual spout contact. We find that inactivating D1 and D2 receptors-expressing medium spiny neurons (MSNs) in the ventrolateral striatum (VLS) oppositely alters the number of licks in a bout. D1- and D2-MSNs exhibit different patterns of lick sequence-related activity and different phases of oscillation time-locked to the lick cycle. On timescale of a lick cycle, transient inactivation of D1-MSNs during tongue protrusion reduces spout contact probability, whereas transiently inactivating D2-MSNs has no effect. On timescale of a lick bout, inactivation of D1-MSNs (D2-MSNs) causes rate increase (decrease) in a subset of basal ganglia output neurons that decrease firing during licking. Our results reveal the distinct roles of D1- and D2-MSNs in regulating licking at both coarse and fine timescales.

## INTRODUCTION

The basal ganglia are critical for movement initiation and execution (Albin et al., 1989; DeLong, 1990; Hikosaka et al., 2000; Klaus et al., 2019; Park et al., 2020). Striatal medium spiny neurons (MSNs) in the input nucleus of the basal ganglia are divided to two classes according to their projection patterns (Gerfen et al., 1990; Gong et al., 2007; Smith et al., 1998). The direct pathway striatal MSNs express the D1-type dopamine receptors (D1-MSNs) and project directly to the substantia nigra pars reticulata (SNr) and internal globus pallidus (GPi), the output nuclei of basal ganglia. The indirect pathway striatal MSNs express the D2-type dopamine receptors (D2-MSNs) and project indirectly to the output nuclei via the external globus pallidus and the subthalamic nucleus. According to the classical rate model, the direct and indirect pathways facilitate and inhibit movement, respectively (Albin et al., 1989; DeLong, 1990). Optogenetic activation of D1-MSNs increases movement, whereas activation of D2-MSNs decreases movement (Kravitz et al., 2010; Oldenburg and Sabatini, 2015), consistent with the rate model. However, cell-type specific recordings found that both D1- and D2-MSNs are activated during voluntary movement (Barbera et al., 2016; Cui et al., 2013; Isomura et al., 2013; Jin et al., 2014; London et al., 2018; Meng et al., 2018), and cell-type specific stimulation showed that activation of either D1- or D2-MSNs can increase or decrease movement velocity (Yttri and Dudman, 2016), inconsistent with the rate model. The concurrent activation of D1- and D2-MSNs is consistent with the support/suppress model, in which both pathways are active to support the desired movement and to suppress the competing movements (Hikosaka et al., 2000; Mink, 1996). Neuronal activity in the direct and indirect pathways also differs in relative timing at sub-second timescales (Markowitz et al., 2018; O’Hare et al., 2016; Sippy et al., 2015) and during different moments of action sequence (Geddes et al., 2018; Jin et al., 2014). It is suggested that the precise activity patterns in both pathways are required for appropriate action sequence (Geddes et al., 2018; Tecuapetla et al., 2016). However, the temporally precise relationship between activity in direct and indirect pathways is not well understood, and the functional role of D1- and D2-MSNs activity at sub-second timescale remains to be investigated.

The outputs of striatal D1- and D2-MSNs converge in the SNr and GPi, whose activity exerts inhibitory influence on neurons in thalamocortical and brainstem motor circuits (DeLong, 1990; Gerfen et al., 1990; Hikosaka et al., 2000; Nelson and Kreitzer, 2014; Redgrave et al., 1999; Smith et al., 1998). The classical model predicts that the direct and indirect pathways inhibit and disinhibit the responses of SNr neurons, respectively (Albin et al., 1989; DeLong, 1990). However, optogenetic activation of D1- or D2-MSNs induced heterogenous response changes in SNr, in which both inhibited and exited cells were observed (Freeze et al., 2013). Optogenetic inhibition of D1-MSNs and D2-MSNs did not result in opposing activity change in SNr neurons as predicted by the classical model (Tecuapetla et al., 2014). During movement, decrease and increase in firings were found for different SNr neurons (Fan et al., 2012; Gulley et al., 2002; Liu et al., 2020), consistent with the proposed function of SNr in the support/suppress model (Mink, 1996). It is unclear how the endogenous activity of D1- or D2-MSNs influences the firings of SNr neurons that exhibit different response patterns during movement execution.

Rhythmic licking behavior in rodent consists of stereotyped tongue movements at a frequency of 6–9 Hz (Rossi et al., 2016; Rossi and Yin, 2015; Weijnen, 1998). In the basal ganglia circuit, the ventrolateral striatum (VLS) receives inputs from motor and somatosensory cortical areas of the mouth region (Hintiryan et al., 2016). Lesions of the VLS in rats impaired tongue movements (Pisa, 1988; Pisa and Schranz, 1988). Activation of direct and indirect pathways in VLS induced and suppressed licking, respectively (Bakhurin et al., 2020; Lee et al., 2020). As both striatal and SNr neurons exhibit lick-related activities (Aldridge and Berridge, 1998; Bakhurin et al., 2016; Gulley et al., 2002; Mittler et al., 1994; Rossi et al., 2016; Sales-Carbonell et al., 2018; Shin et al., 2018), it is of interest to use licking behavior to examine the temporal response patterns of D1- and D2-MSNs in VLS, and how striatal activities in the two pathways regulate the responses of SNr neurons during licking.

In this study, we used a voluntary licking task in head-fixed mice to examine the roles of D1- and D2-MSNs in regulating licking movement. On a fine timescale within hundreds of milliseconds, we found that D1- and D2-MSNs exhibited distinct patterns of oscillatory activity during the lick cycle, and transient inactivation of D1- but not D2-MSNs reduced the probability of spout contact. During execution of licking, inactivation of D1- and D2-MSNs led to distinct response changes in sub-populations of lick-related SNr neurons. Thus, our data support that the direct and indirect pathway striatal neurons exhibit temporally distinct activity patterns and differentially modulates basal ganglia output responses.

## RESULTS

### Opponent regulation of licking behavior by D1- and D2-MSNs in VLS

Head-fixed mice were trained to voluntarily lick from a drinking spout. Tongue licks were detected using an electrical or infrared lick sensor. The spout contact duration of an individual lick was defined as the time period the tongue contacted the spout, whereas the inter-contact interval was the interval between the onsets of two consecutive spout contacts (Figure S1A). To encourage the mice to display clusters of licks as well as periods of no licks (Figure 1A), the delivery of sucrose (10%) was triggered if a spout contact was preceded by at least 1 s of no contacts (Figure S1B). A valid lick bout, which triggered the delivery of sucrose and the subsequent consumption, was defined as a group of licks in which the first spout contact was preceded by ≥ 1 s of no contacts, the inter-contact interval of the first three contacts was ≤ 0.3 s, and the last contact was followed by ≥ 0.5 s of no contacts. We found that the spout contact duration was 40.53 ± 1.66 ms (mean ± SEM) and the inter-contact interval was 146.24 ± 1.99 ms (mean ± SEM, n = 9 mice, Figure S1C), consistent with the licking frequency of 6–9 Hz reported by previous studies (Rossi and Yin, 2015; Weijnen, 1998). When the time window of sucrose delivery lasted for 4 s, the number of licks in a bout was 23 ± 1.66 (mean ± SEM), and the bout length was 3.41 ± 0.22 s (mean ± SEM, n = 9 mice, Figure S1C).

**Figure 1.**
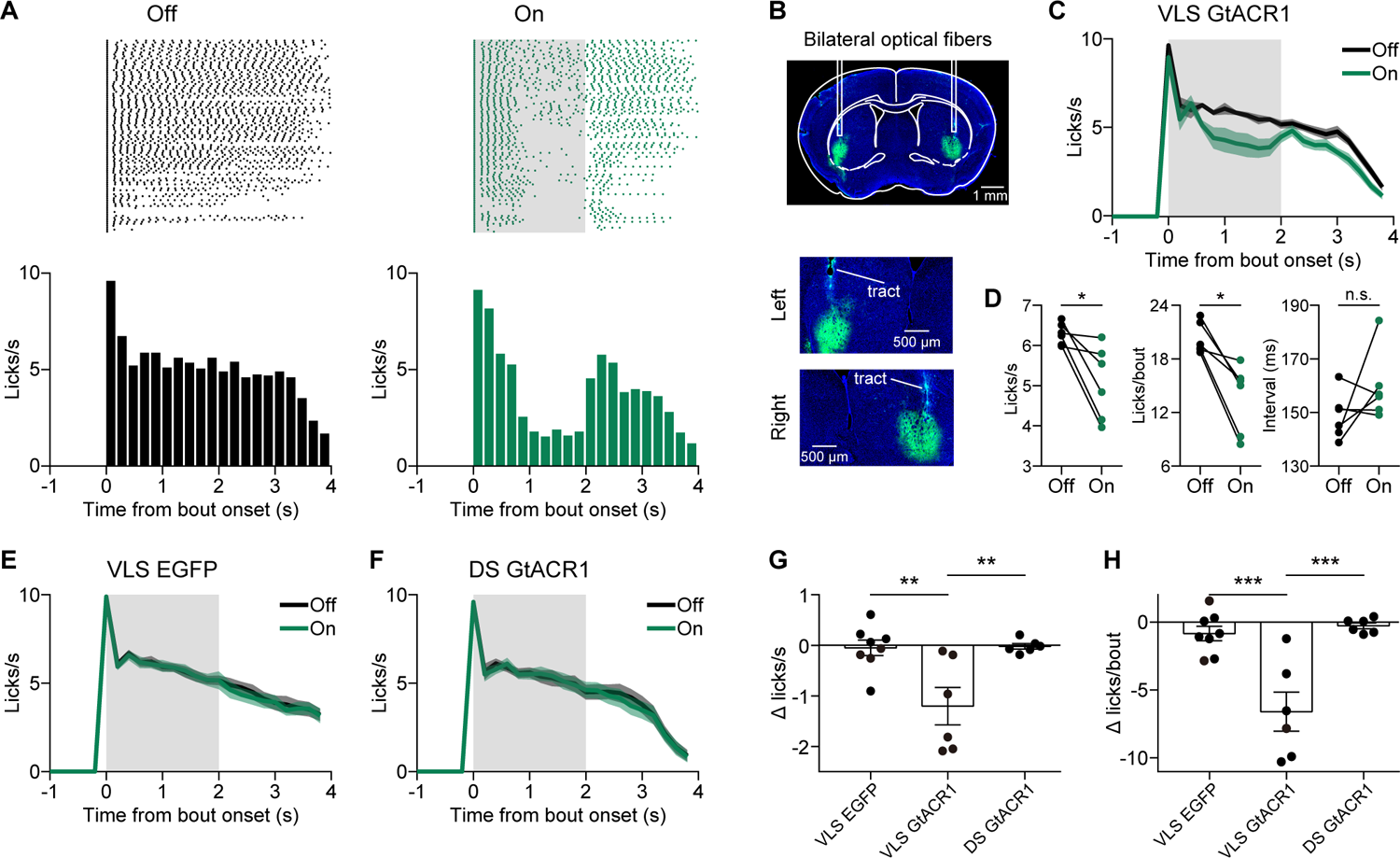
VLS is involved in motor control of licking behavior. (A) Lick rasters and lick PSTHs for laser-off and laser-on trials of an example mouse, in which AAV-GtACR1 was bilaterally injected in VLS. (B) Representative fluorescence image showing the expression of AAV-hSyn-hGtACR1-EGFP in VLS of a C57BL/6 mouse. Lower, enlarged views showing fiber tracts. (C) Inactivation of VLS significantly decreased the amplitude of lick PSTH (F_(1, 5)_ = 10.51, p = 0.023, two-way repeated measures ANOVA). (D) Inactivation of VLS significantly reduced lick rate and number of licks per bout, without affecting inter-contact interval. * p < 0.05, n = 6, Wilcoxon signed rank test. (E) Laser stimulation did not change the lick PSTH in control mice, in which AAV-EGFP was bilaterally injected in VLS. (F) Inactivation of DS did not change the lick PSTH. (G) Comparison of the change in lick rate (between laser-on and laser-off trials) among the three groups of mice (VLS-EGFP: n = 8; VLS-GtACR1: n = 6; DS-GtACR1: n = 6). ** p < 0.01, one-way ANOVA followed by Sidak’s multiple comparisons test. (H) Comparison of the change in number of licks per bout (between laser-on and laser-off trials) among the three groups of mice (VLS-EGFP: n = 8; VLS-GtACR1: n = 6; DS-GtACR1: n = 6). *** p < 0.001, one-way ANOVA followed by Sidak’s multiple comparisons test. Data represent mean ± SEM. Gray shading, duration of laser stimulation. See also Figure S1 and S2.

To examine the necessity of VLS activity in regulating licking action, we bilaterally injected AAV2/5-hSyn-hGtACR1-EGFP-WPRE in VLS and implanted optical fibers above the injection site (Figure 1B). In laser-on trials, 2 s of laser stimulation was triggered by the first spout contact of a bout that was preceded by ≥ 1 s of no contacts (Figure S1B and Figure 1A). For both laser-off and laser-on trials, the time window of sucrose delivery lasted for 3 s. In vivo recordings verified that laser stimulation reduced the spontaneous firing rates of VLS neurons in awake mice (Figure S2A and S2B). Across the population of mice, optogenetic inactivation of VLS caused a significant reduction in the amplitude of lick PSTH (F_(1, 5)_ = 10.51, p = 0.023, two-way repeated measures ANOVA, Figure 1C). We computed lick rate using licks within the first 2 s of lick bout, and found that optogenetic inactivation of VLS resulted in a significant decrease of lick rate (p = 0.031, n = 6, Wilcoxon signed rank test, Figure 1D). Inactivation of VLS reduced the number of licks per bout (p = 0.031), without affecting the inter-contact interval (p = 0.44, n = 6, Wilcoxon signed rank test, Figure 1D), suggesting that VLS inactivation causes reduction in lick rate by early termination of lick bout without affecting the lick rhythm. As the number of licks in a bout may decrease over trials, we further examined the effect of laser stimulation by dividing trials in the same session into three blocks, which contained equal (or nearly equal) number of trials (Figure S1D and S1E). We found that the laser-induced reduction in lick rate or in number of licks per bout did not differ significantly across the three blocks (Δ licks/s: F_(1.30, 6.51)_ = 0.6, p = 0.51; Δ licks/bout: F_(1.61, 8.04)_ = 0.1, p = 0.86, one-way repeated measures ANOVA, Figure S1D and S1E). For control mice in which AAV2/8-hSyn-eGFP-3Flag-WPRE-SV40pA was injected in VLS (Figure S1F), laser stimulation did not affect the lick PSTH (F_(1, 7)_ = 0.097, p = 0.77, two-way repeated measures ANOVA, Figure 1E).

Unlike VLS inactivation, inactivation of the dorsal striatum (DS) did not cause a significant change in the amplitude of lick PSTH (F_(1, 5)_ = 0.11, p = 0.75, two-way repeated measures ANOVA, Figure 1F and Figure S1G). Among the three groups of mice (VLS-EGFP, VLS-GtACR1 and DS-GtACR1), the reduction in lick rate and in number of licks per bout for the VLS-GtACR1 group was significantly larger than those for other groups (Δ licks/s: F_(2, 17)_ = 8.71, p = 2.48×10^-3^, Δ licks/bout: F_(2, 17)_ = 16.23, p = 1.14×10^-4^, one-way ANOVA followed by Sidak’s multiple comparisons test, p < 0.01, Figure 1G and 1H). These results demonstrate that the VLS but not the DS is involved in motor control of licking behavior.

To test the role of D1- and D2-MSNs in licking behavior, we bilaterally injected AAV2/8-CAG-DIO-GtACR1-P2A-EGFP in the VLS of D1-Cre or D2-Cre mice (Figure 2A and 2E). Slice recordings confirmed that laser stimulation could suppress the firing rates of D1- and D2-MSNs in VLS (Figure S2C). We found that inactivation of D1-MSNs with 2-s laser stimulation significantly decreased the amplitude of lick PSTH (F_(1, 16)_ = 20.77, p = 3.23×10^-4^, two-way repeated measures ANOVA), lick rate (p = 4.58×10^-5^) and number of licks per bout (p = 5.34×10^-4^, n = 17 mice, Wilcoxon signed rank test, Figure 2B−2D), whereas inactivation of D2-MSNs significantly increased the amplitude of lick PSTH (F_(1, 12)_ = 36.41, p = 5.9×10^-5^, two-way repeated measures ANOVA), lick rate (p = 2.44×10^-4^) and number of licks per bout (p = 1.46×10^-3^, n = 13 mice, Wilcoxon signed rank test, Figure 2F−2H). Inactivation of D1-MSNs or D2-MSNs did not affect the inter-contact interval (p = 0.93 and 0.54, respectively, Wilcoxon signed rank test, Figure 2D and 2H). When we divided trials in the same session into three blocks, we found that the effect of inactivating D1-MSNs was stronger in the first block (Figure S3A and S3B), whereas the effect of inactivating D2-MSNs on licking (Figure S3C and S3D), particularly for the number of licks per bout, was stronger in the last block. Laser stimulation did not affect lick PSTH, lick rate or number of licks per bout for control D1-Cre or D2-Cre mice in which AAV2/8-hSyn-FLEX-tdTomato was injected in the VLS (Figure S3E–S3G). Thus, the activity of D1- and D2-MSNs in VLS oppositely regulates licking action, consistent with a previous report (Bakhurin et al., 2020).

**Figure 2.**
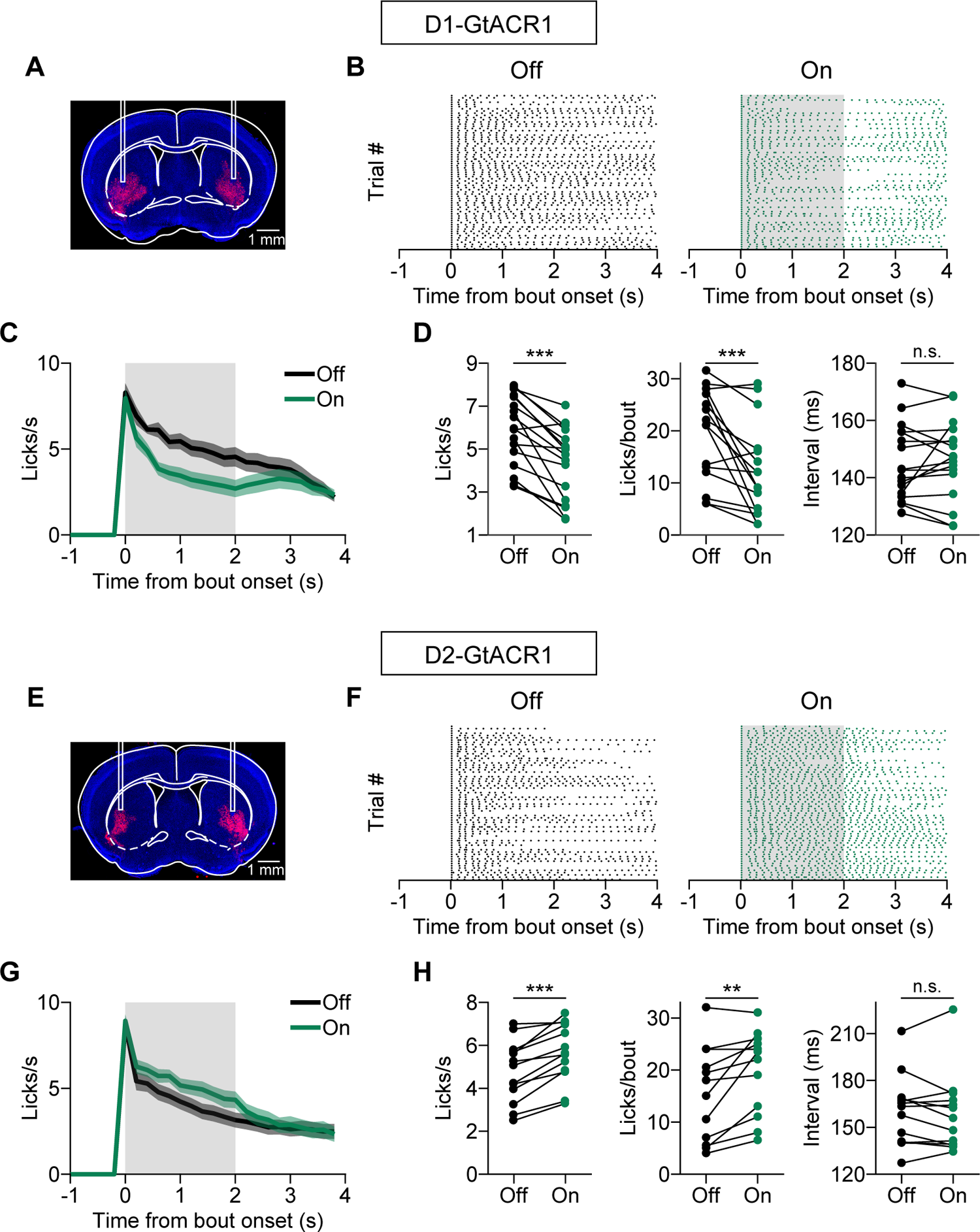
Opponent regulation of licking behavior by D1- and D2-MSNs in VLS. (A) Representative fluorescence image showing the expression of AAV-DIO-GtACR1-EGFP in VLS of a D1-Cre mouse. The EGFP was observed using immunohistochemistry. (B) Lick rasters in laser-off and laser-on trials for an example D1-Cre mouse, in which AAV-DIO-GtACR1 was bilaterally injected in VLS. (C) Inactivation of D1-MSNs significantly decreased the amplitude of lick PSTH (F_(1, 16)_ = 20.77, p = 3.23×10^-4^, two-way repeated measures ANOVA). (D) Inactivation of D1-MSNs significantly decreased lick rate and number of licks per bout, without affecting inter-contact interval. *** p < 0.001, n = 17 mice, Wilcoxon signed rank test. (E) Representative fluorescence image showing the expression of AAV-DIO-GtACR1-EGFP in VLS of a D2-Cre mouse. The EGFP was observed using immunohistochemistry. (F) Lick rasters in laser-off and laser-on trials for an example D2-Cre mouse, in which AAV-DIO-GtACR1 was bilaterally injected in VLS. (G) Inactivation of D2-MSNs significantly increased the amplitude of lick PSTH (F_(1, 12)_ = 36.41, p = 5.9×10^-5^, two-way repeated measures ANOVA). (H) Inactivation of D2-MSNs significantly increased lick rate and number of licks per bout, without affecting inter-contact interval. ** p < 0.01, *** p < 0.001, n = 13 mice, Wilcoxon signed rank test. Data represent mean ± SEM. Gray shading, duration of laser stimulation. See also Figure S2 and S3.

### D1- and D2-MSNs display different lick-sequence related activity

To examine the activity patterns of D1- and D2-MSNs in VLS, we performed electrophysiological recordings using optrode from D1-Cre or D2-Cre mice in which AAV2/8-Syn-FLEX-ChrimsonR-tdTomato were injected in VLS (Figure 3A, 3C and Figure S4A). A unit was identified as ChrimsonR-expressing if it satisfied the two criteria (Lee et al., 2019): 1) significantly responded to laser stimulation with a spike latency < 6 ms, and 2) the Pearson’s correlation coefficient between waveforms of spontaneous spikes and laser-evoked spikes was > 0.95 (Figure 3B, 3D and Figure S4B). Licking parameters, including inter-contact interval, number of licks per bout and bout length, were similar between D1-Cre and D2-Cre mice (Figure S4C). We classified the identified ChrimsonR-expressing units into putative MSNs and other neurons according to their spike waveforms and coefficients of variation (CV) of inter-spike intervals (Shin et al., 2018) (Figure S4D and S4E). The responses of MSNs (Figure 3E-3H) during licking behavior were analyzed.

**Figure 3.**
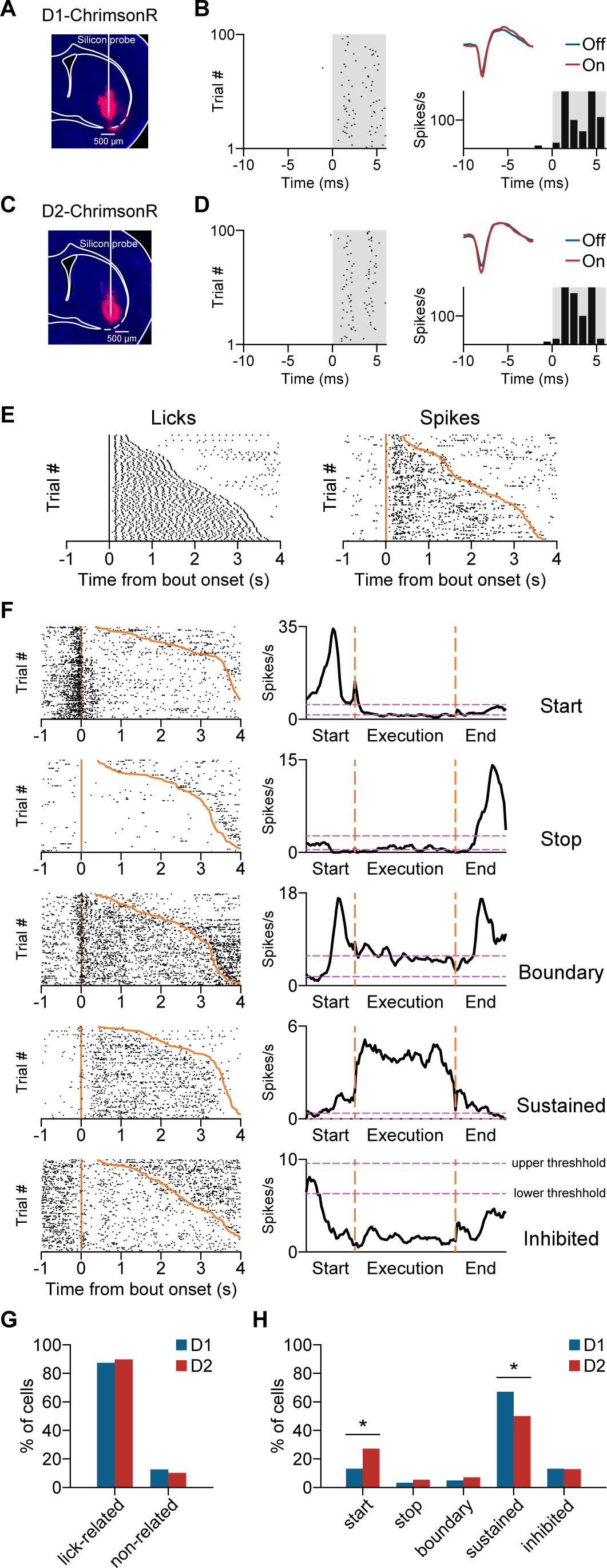
D1- and D2-MSNs display different lick-sequence related activity. (A) Fluorescence image of VLS for an example D1-Cre mouse in which AAV-FLEX-ChrimsonR-tdTomato was injected in VLS. (B) Spike rasters (left), waveforms and PSTH (right) of an identified D1-MSN. Time 0 is laser onset. Gray shading, laser stimulation. (C) Fluorescence image of VLS for an example D2-Cre mouse in which AAV-FLEX-ChrimsonR-tdTomato was injected in VLS. (D) Spike rasters, waveforms and PSTH of an identified D2-MSN. Similar to that described in (B). (E) Lick rasters (left) and spike rasters (right), sorted by the length of lick bout, for an identified D1-MSN during licking behavior. Orange line and curve: start and end of lick bout. (F) Spike rasters (left) and PSTHs (right, smoothed with a moving average of consecutive 5 bins) for example MSNs that were classified as start, stop, boundary, sustained, and inhibited neurons. In the spike rasters, orange line and curve indicate the start and end of lick bout, respectively. In the PSTHs, the two vertical dashed lines denote the start and end of the execution phase. (G) The percentages of lick-related and non-related neurons for D1-MSNs (n = 87) and D2-MSNs (n = 78). (H) Lick-related D1- and D2-MSNs differed in the percentage of start neurons and sustained neurons. * p < 0.05, Fisher’s exact test. See also Figure S4.

Previous studies using lever-press task found that striatal neurons signal the initiation, execution, and termination of learned action sequence (Jin and Costa, 2010, 2015; Jin et al., 2014; Martiros et al., 2018; Vandaele et al., 2019). We examined whether the activities of D1- and D2-MSNs in VLS are related to the execution or start/stop of licking. A trial of lick bout was defined to include a baseline phase (from - 1 s to −0.5 s relative to bout onset), a start phase (from −0.5 s to 0.25 s relative to bout onset), an execution phase (the duration of the lick bout), and a stop phase (from −0.25 s to 0.5 s relative to bout offset). We computed the mean and SD of firing rate in baseline phase, and defined an upper and a lower threshold as 3 SD above and below the mean, respectively (horizontal, purple dashed lines in Figure 3F, right panels). By comparing the responses in the execution (start, or stop) phase with the upper (or lower) threshold, we classified a unit as start, stop, boundary, sustained, or inhibited neuron (Figure 3F, see STAR Methods). Neurons that were classified as one of these response types were considered to exhibit lick-related activities. We found that the percentage of lick-related neurons did not differ between D1- and D2-MSNs (χ^2^_(1)_ = 0.23, p = 0.63, n = 87 and 78 for D1- and D2-MSNs, respectively, Figure 3G). Among the lick-related neurons, the percentage of start neurons was significantly higher in D2-MSNs than in D1-MSNs (p = 0.04, Fisher’s exact test), that of sustained neurons was significantly higher in D1-MSNs than in D2-MSNs (p = 0.044, Fisher’s exact test), and that of stop (boundary or inhibited) neurons was not significantly different between D1- and D2-MSNs (p > 0.5, Fisher’s exact test, Figure 3H). These results suggest that, while D1- and D2-MSNs are both activated during licking movement, they have different activity during the initiation and execution of lick sequence.

### D1- and D2-MSNs exhibit different phase relationships with the lick cycle

A prominent feature of licking behavior is the rhythmic alternation of tongue protrusions and retractions (Travers et al., 1997; Weijnen, 1998). The period of a lick cycle was ∼150 ms, which could be observed from the peri-event time histogram (PETH) computed using licks around ±200 ms of each spout-contact (Figure 4A−4D, left panels) (Rossi et al., 2016). We next examined whether MSNs in VLS show oscillatory activity related to the lick cycle. We calculated spout-contact-triggered spike PETH using spikes occurring around ±200 ms of each contact (Figure 4A−4D, middle panels), and then computed the Pearson’s correlation coefficient between the lick PETH and the spout-contact-triggered spike PETH (Figure 4A−4D, right panels) or between the lick PETH and the spike PETH shifted by 40 ms (corresponding to a phase shift of ∼90°) (Figure S5). For those neurons in which the correlation coefficient was significant, the spout-contact-triggered spike PETHs showed oscillatory activity (Figure 4A−4D and Figure S5). Among the optogenetically identified MSNs, the percentages of D1- and D2-MSNs showing lick-related oscillatory activity were 75.9% (66 out of 87 units) and 74.4% (58 out of 78 units), respectively. To determine the oscillatory phase relative to the lick cycle (−180° to 180°), we identified the time at which the amplitude of spike PETH was maximum and transformed it to polar coordinate. The distribution of phases for D1-MSNs was significantly different from a uniform distribution (p = 2.36×10^-6^, n = 66, Rayleigh test, Figure 4E), whereas that for D2-MSNs was not (p = 0.33, n = 58, Rayleigh test, Figure 4F). The phase of peak response was significantly different between D1- and D2-MSNs (p = 5.49×10^-4^, circular Watson-Williams two-sample test (Berens, 2009)), and the z-scored spike PETHs averaged across oscillatory units were significantly different between D1- and D2-MSNs (F_interaction(35, 4270)_ = 7.76, p < 1×10^-15^, two-way ANOVA with mixed design, Figure 4G). Thus, the activity of D1- and D2-MSNs exhibited distinct phase relationships with the lick cycle. Interestingly, the peak response of population D1-MSNs with oscillatory activity was around 0° (Figure 4G), corresponding to the time of spout contact.

**Figure 4.**
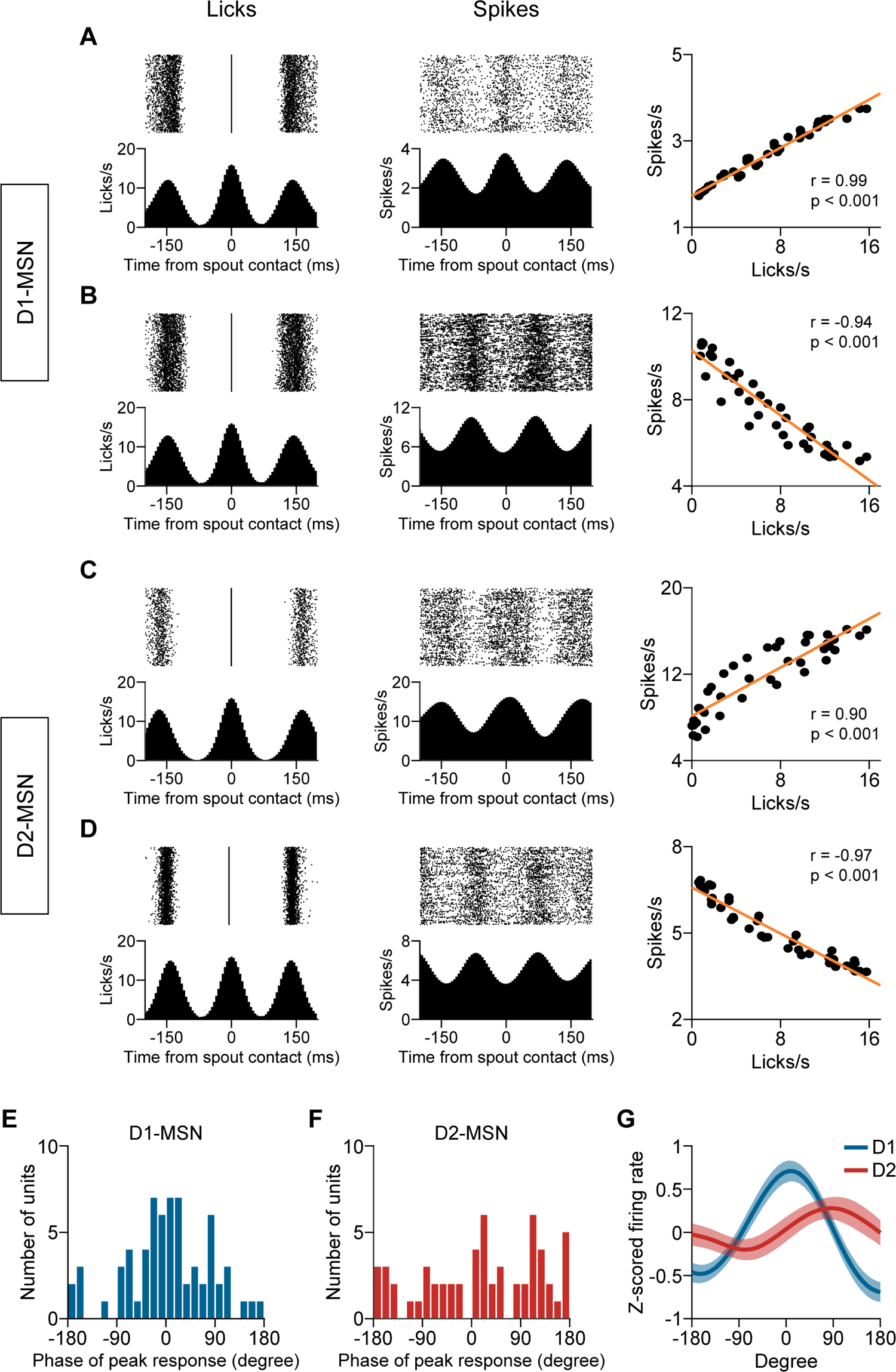
D1- and D2-MSNs exhibit different phase relationships with the lick cycle. (A) Left, lick rasters and lick PETH for an example D1-Cre mouse. Middle, rasters and PETH of spout-contact-triggered spikes for a D1-MSN recorded from this mouse. Right, correlation between lick PETH and spout-contact-triggered spike PETH. (B) Similar to that described in (A) for another D1-MSN. (C) Similar to that described in (A) except that the licks (spikes) were from an example D2-Cre mouse (an example D2-MSN). (D) Similar to that described in (C) for another D2-MSN. (E) Distribution of the phase at the peak of oscillatory activity for D1-MSNs (n = 66). (F) Distribution of the phase at the peak of oscillatory activity for D2-MSNs (n = 58). (G) Population average (mean ± SEM) of z-scored spout-contact-triggered spike PETH for D1-MSNs and D2-MSNs (F_interaction(35, 4270)_ = 7.76, p < 1×10^-15^, two-way ANOVA with mixed design). See also Figure S5.

### Transient inactivation of D1-MSNs during tongue protrusion reduces spout-contact probability

We wondered whether transient perturbation of D1- or D2-MSNs activity at a timescale within the lick cycle affects individual spout-contact. We bilaterally injected AAV2/8-CAG-DIO-GtACR1-P2A-EGFP in the VLS of D1-Cre or D2-Cre mice, which were then trained to perform voluntary licking behavior. As the phase of peak oscillatory activity in many D1-MSNs was around 0°, we applied transient laser stimulation (duration = 25 ms) around the time of tongue protrusion. After the first spout-contact, laser stimulation was triggered after an inter-contact interval following the onset of the second contact (Figure 5A and 5D). Slice recordings showed that 25-ms laser stimulation rapidly suppressed the firing rates of D1- and D2-MSNs, whose activity gradually recovered starting from ∼150 ms after laser onset (Figure S6A and S6B). We found that transient inactivation of D1-MSNs significantly reduced the amplitude of spout contact signal for the third contact (F_(1, 6)_ = 78.2, p = 1.16×10^-4^, two-way repeated measures ANOVA, Figure 5B) and the probability of the third contact (p = 0.016, Wilcoxon signed rank test), without affecting the probability of the fourth contact (p = 0.58) or the length of lick bout (p = 0.58, n = 7 mice, Wilcoxon signed rank test, Figure 5C). When we divided trials in the same session into 3 blocks, we found that the reduction in the probability of third spout-contact did not differ across blocks (Figure S6C). For the experiments in which D1-MSNs were inactivated with 2-s laser stimulation (Figure 2C), we observed a smaller reduction in the spout-contact probability immediately after laser onset (Figure S6D), likely due to a lower laser power compared with that of 25-ms laser stimulation. When we applied the same power as that in 25-ms laser stimulation, we found that 2-s inactivation of D1-MSNs greatly reduced the spout-contact probability immediately after laser onset (Figure S6E). Unlike perturbation of D1-MSNs, transient inactivation of D2-MSNs with 25-ms laser stimulation around the time of tongue protrusion did not change the spout contact signal (F_(1, 5)_ = 1.2, p = 0.32, two-way repeated measures ANOVA), the probability of the third contact (p = 0.69) or the length of lick bout (p = 1, n = 6 mice, Wilcoxon signed rank test, Figure 5E and 5F). Inactivation of D2-MSNs for 150 ms at a later time (3 s after first spout-contact), when lick rate was lower, still did not affect licking behavior (Figure S6F). For control D1-Cre mice in which AAV2/8-hSyn-FLEX-tdTomato was injected in VLS, the amplitude of spout contact signal, the probability of spout-contact or bout length was not affected by transient laser stimulation (Figure S6G−S6I). Thus, the activity of D1-MSNs in VLS around tongue protrusion is necessary for the execution of individual spout-contact.

**Figure 5.**
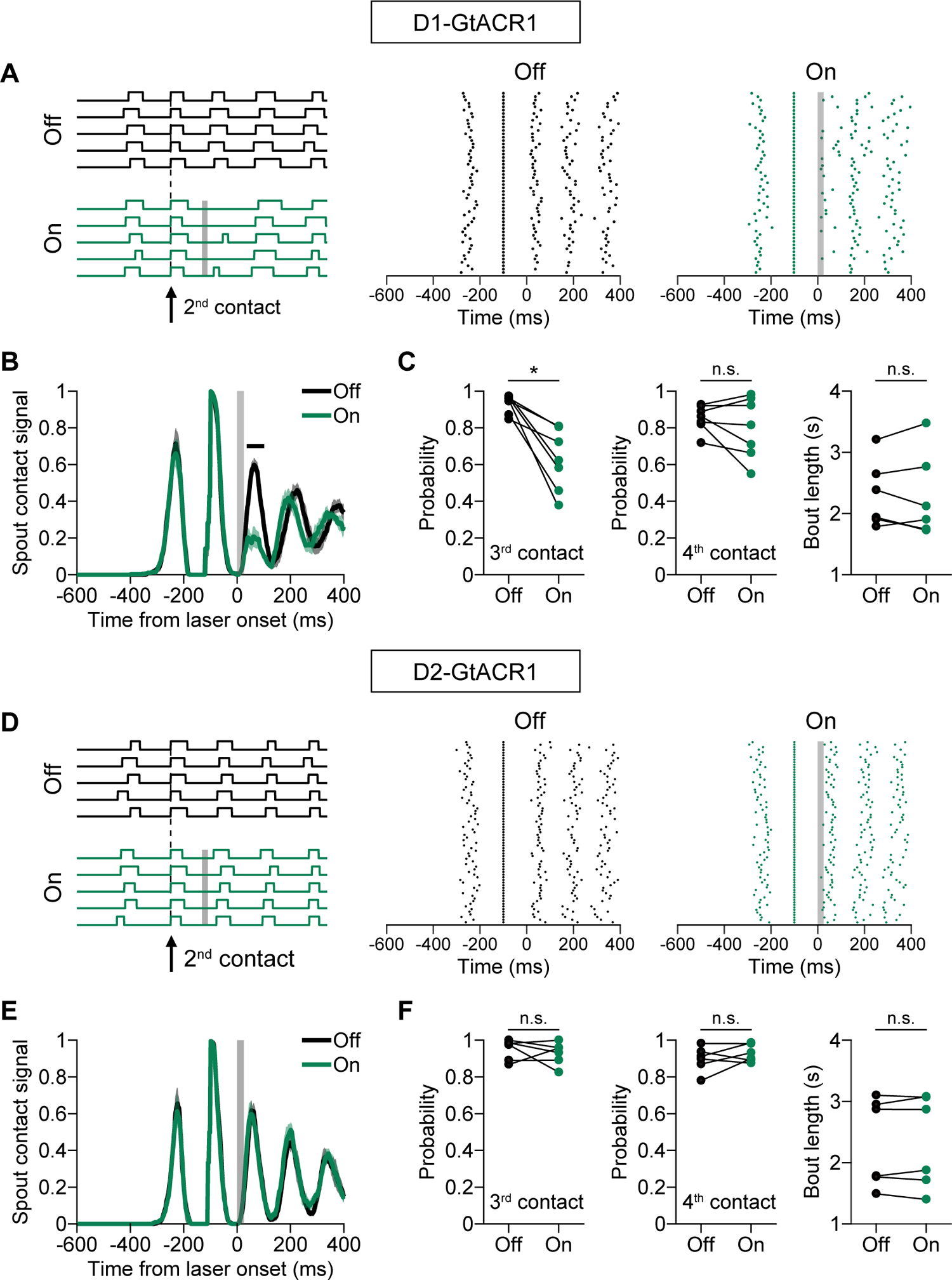
Transient inactivation of D1-MSNs during tongue protrusion reduces spout contact probability. (A) Licking behavior for an example D1-Cre mouse in which AAV-DIO-GtACR1 was bilaterally injected in VLS. Left, example spout contact signals (five laser-off and five laser-on trials). The signals were detected by the interruption of an infrared beam and sign-reversed for display purpose (aligned to the onset of second spout-contact). Right, lick rasters of all trials. Time 0 is laser onset. Gray shading, duration of laser stimulation. (B) Transient inactivation of D1-MSNs around the time of tongue protrusion of the third spout-contact significantly reduced the amplitude of spout contact signal (F_(1, 6)_ = 78.2, p = 1.16×10^-4^, two-way repeated measures ANOVA). Black horizontal line marks the time points in which the signal amplitude was significantly different between laser-off and laser-on conditions. Gray shading, duration of laser stimulation. Data represent mean ± SEM. (C) Transient inactivation of D1-MSNs around the time of tongue protrusion significantly reduced the probability of the third spout-contact, without affecting the probability of the fourth spout-contact or the bout length. * p < 0.05, n = 7 mice, Wilcoxon signed rank test. (D) Spout contact signals and lick rasters for an example D2-Cre mouse in which AAV-DIO-GtACR1 was bilaterally injected in VLS. Similar to that described in (A). (E) Transient inactivation of D2-MSNs around the time of tongue protrusion of the third spout-contact did not change the spout contact signal. Similar to that described in (B). (F) Transient inactivation of D2-MSNs around the time of tongue protrusion did not change spout contact probability or bout length. n = 6 mice, Wilcoxon signed rank test. See also Figure S6.

### Differential roles of endogenous D1- and D2-MSNs activity in controlling SNr responses during licking

By injecting CTB in VLS, we observed anterogradely labelled fibers in the lateral SNr (Figure S7A), consistent with previous reports (Lee et al., 2020; von Krosigk et al., 1992). To examine how the endogenous activity of D1- or D2-MSNs influences the downstream neurons, we bilaterally injected AAV2/8-CAG-DIO-GtACR1-P2A-EGFP in VLS of D1-Cre or D2-Cre mice. We performed extracellular recordings from the lateral SNr of licking mice, with and without laser stimulation in VLS (Figure 6A, 6B, 7A, and 7B). The units were classified as GABAergic and non-GABAergic neurons based on their spike waveforms (Figure S7B), and GABAergic SNr neurons were used for subsequent analysis.

**Figure 6.**
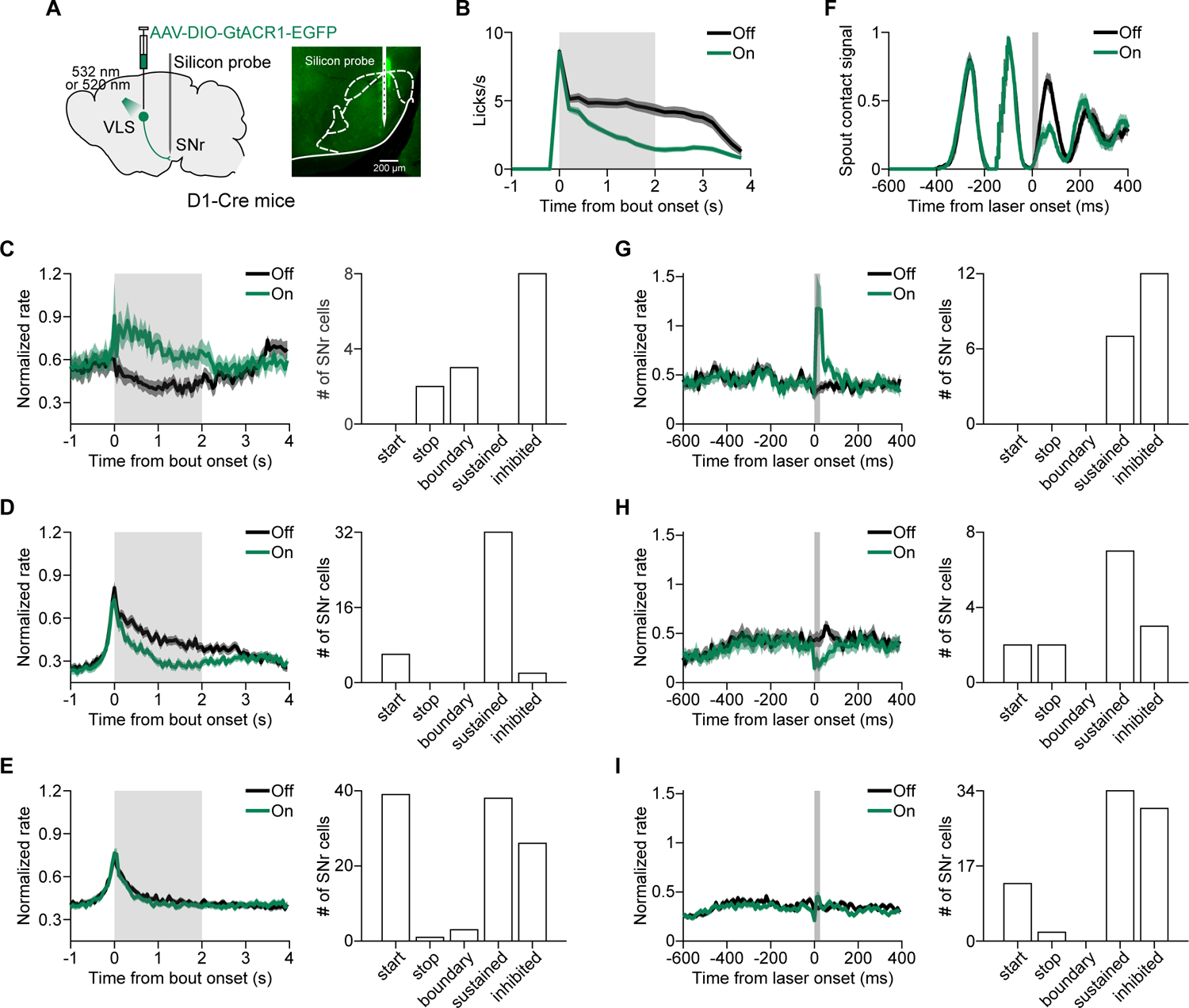
Effect of inactivating D1-MSNs in VLS on SNr responses during licking. (A) Left, schematic of recording from SNr while bilateral inactivation of D1-MSNs in VLS. Right, example fluorescence image showing the electrode track in SNr marked by DiO. (B) For mice used in SNr recordings, inactivation of D1-MSNs significantly reduced the amplitude of lick PSTH (F_(1, 18)_ = 44.46, p = 2.96×10^-6^, two-way repeated measures ANOVA). (C) Left, the responses of SNr neurons (n = 13) that showed significant increase in firing rates after inactivation of D1-MSNs. For each neuron, the responses were normalized by the peak firing rate in laser-off condition. Right, distribution of response types for those SNr neurons that significantly increased firing rate after inactivation of D1-MSNs. (D) Left, the responses of SNr neurons (n = 40) that showed significant decrease in firing rates after inactivation of D1-MSNs. Similar to that described in (C). Right, distribution of response types for those SNr neurons that significantly decreased firing rate. (E) Left, the responses of SNr neurons (n = 107) that did not significantly change firing rates after inactivation of D1-MSNs. Similar to that described in (C). Right, distribution of response types for those SNr neurons that did not significantly change firing rate. (F) For D1-Cre mice used in SNr recordings, transient inactivation of D1-MSNs around the time of tongue protrusion significantly reduced the amplitude of spout contact signal for the third contact (F _(1, 11)_ = 40.8, p = 5.2×10^-5^, two-way repeated measures ANOVA). (G) Left, the responses of SNr neurons (n = 19) that showed significant rate increase within the time window of the third contact after transient inactivation of D1-MSNs. For each neuron, the responses were normalized by the peak firing rate in laser-off condition. Right, distribution of response types for those SNr neurons that significantly increased firing rate within the time window of the third contact after transient inactivation of D1-MSNs. (H) Left, the responses of SNr neurons (n = 14) that showed significant rate decrease within the time window of the third contact after transient inactivation of D1-MSNs. Similar to that described in (G). Right, distribution of response types for those SNr neurons that significantly decreased firing rate within the time window of the third contact. (I) Left, the responses of SNr neurons (n = 79) that did not significantly change firing rate within the time window of the third contact after transient inactivation of D1-MSNs. Similar to that described in (G). Right, distribution of response types for those SNr neurons that did not significantly change firing rate within the time window of the third contact. Data represent mean ± SEM. Gray shading, duration of laser stimulation. See also Figure S7.

For D1-Cre mice, we performed two sets of experiments. In the first set of experiment, 2-s laser stimulation was triggered by the first spout-contact of a bout (Figure S7C). Inactivation of D1-MSNs in VLS caused a significant decrease in the amplitude of lick PSTH (F_(1, 18)_ = 44.46, p = 2.96×10^-6^, two-way repeated measures ANOVA, Figure 6B), and both increase and decrease in firing rates of SNr neurons were observed (Figure S7F). At the population level, inactivation of D1-MSNs significantly increased the firing rates in 8.13% (13/160) of SNr neurons, most of which were classified as inhibited neurons in laser-off condition (Figure 6C), and significantly decreased the firing rates in 25% (40/160) of SNr neurons, which were predominantly sustained neurons (Figure 6D). For a large fraction of SNr neurons (66.87%, 107/160), the firing rates were not significantly affected by inactivation of D1-MSNs (Figure 6E). In the second set of experiment, 25-ms laser stimulation was applied around the time of tongue protrusion for the third spout-contact in a bout (Figure S7D), leading to a significant decrease in the amplitude of spout contact signal of the third contact (F _(1, 11)_ = 40.8, p = 5.2×10^-5^, two-way repeated measures ANOVA, Figure 6F). When we analyzed responses within time window of the third contact, we found firing rate increase in 16.96% (19/112) of SNr neurons, which were predominantly inhibited neurons (Figure 6G), and firing rate decrease in 12.5% (14/112) of SNr neurons, most of which were sustained neurons (Figure 6H). Similar to the effect of 2-s laser stimulation, the responses of most SNr neurons (70.54%, 79/112) were not significantly affected by transient inactivation of D1-MSNs (Figure 6I). Thus, on both timescales of seconds and sub-seconds, inactivation of D1-MSNs resulted in rate increase (decrease) for a subset of SNr neurons that showed inhibited (sustained) responses during licking. For D2-Cre mice, we examined the effect of 2-s laser stimulation in VLS (Figure S7E), which significantly increased the amplitude of lick PSTH (F _(1, 16)_ = 69.26, p = 3.31×10^-7^, two-way repeated measures ANOVA, Figure 7B), on the responses of SNr neurons (Figure S7G). In contrast to that observed in D1-Cre mice, inactivation of D2-MSNs in VLS significantly decreased the firing rates in 7.77% (8/103) of SNr neurons that were mostly of the inhibited response type (Figure 7C), and significantly increased the firing rates in 21.36% (22/103) of SNr neurons that were predominantly sustained neurons (Figure 7D). Firing rates in 70.87% (73/103) of SNr neurons were not affected by inactivation of D2-MSNs (Figure 7E).

**Figure 7.**
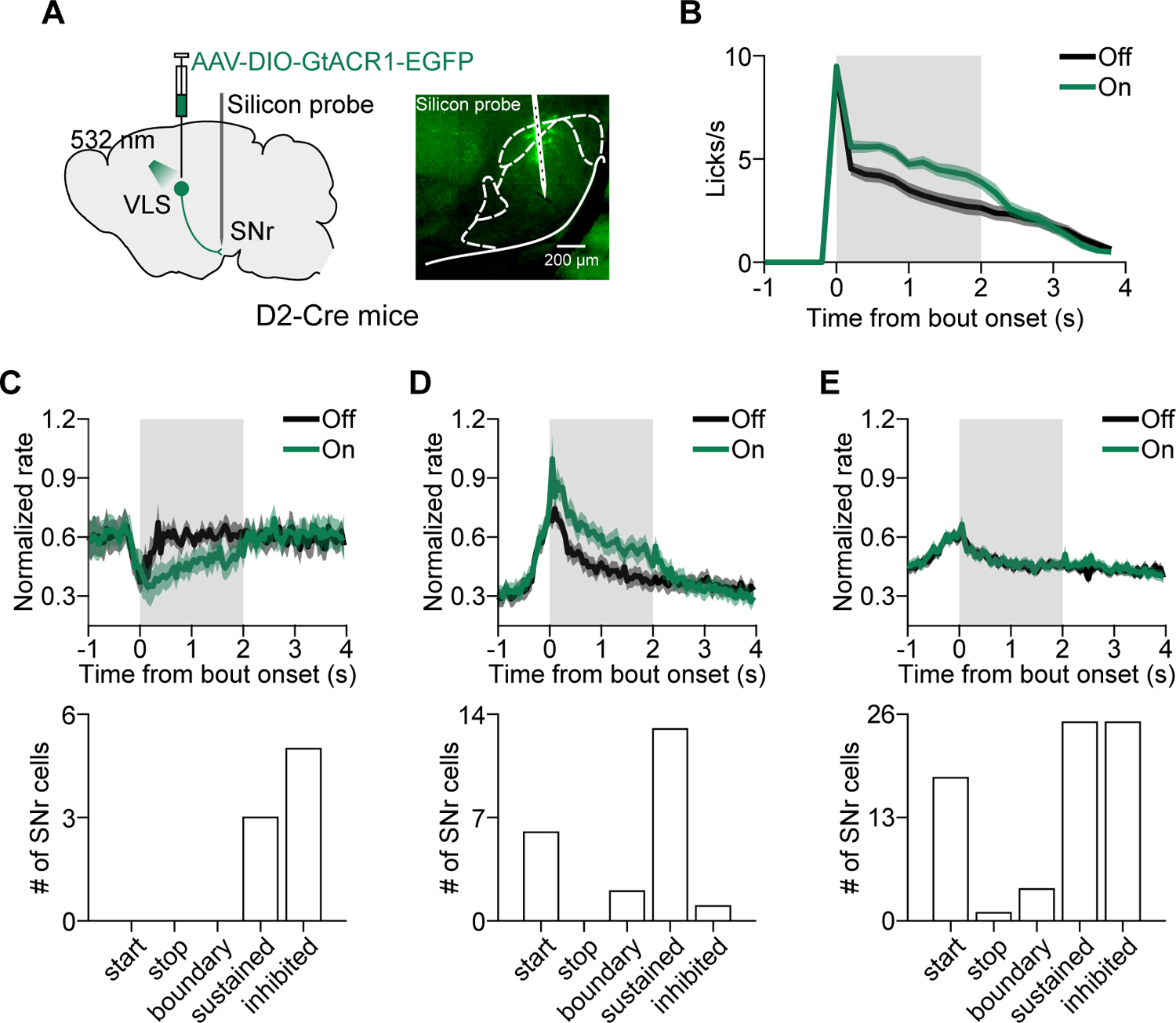
Effect of inactivating D2-MSNs in VLS on SNr responses during licking. (A) Left, schematic of recording from SNr while bilateral inactivation of D2-MSNs in VLS. Right, example fluorescence image showing the electrode track in SNr marked by DiO. (B) For D2-Cre mice used in SNr recordings, inactivation of D2-MSNs significantly increased the amplitude of lick PSTH (F _(1, 16)_ = 69.26, p = 3.31×10^-7^, two-way repeated measures ANOVA). (C) Upper, the responses of SNr neurons (n = 8) that showed significant decrease in firing rates after inactivation of D2-MSNs. For each neuron, the responses were normalized by the peak firing rate in laser-off condition. Lower, distribution of response types for those SNr neurons that significantly decreased firing rate after inactivation of D2-MSNs. (D) Upper, the responses of SNr neurons (n = 22) that showed significant increase in firing rates after inactivation of D2-MSNs in. Similar to that described in (C). Lower, distribution of response types for those SNr neurons that significantly increased firing rate. (E) Upper, the responses of SNr neurons (n = 73) that did not significantly change firing rates after inactivation of D2-MSNs. Similar to that described in (C). Lower, distribution of response types for those SNr neurons that did not significantly change firing rate. Data represent mean ± SEM. Gray shading, duration of laser stimulation. See also Figure S7.

Thus, the results demonstrated that the endogenous activity of D1- and D2-MSNs oppositely regulates the responses of a subset of SNr neurons during licking.

## DISCUSSION

Using voluntary licking behavior in mice, we found that D1- and D2-MSNs in VLS played distinct roles in regulating licking movement. Optogenetic tagging showed that D1- and D2-MSNs exhibited different patterns of lick sequence-related activity. At a fine timescale, they also differed in the oscillatory activity time-locked to the lick cycle. Transient inactivation of D1-MSNs around the time of tongue protrusion reduced the spout-contact probability, whereas transient inactivation of D2-MSNs did not affect licking. Inactivation of D1- and D2-MSNs produced opposing effects in a subset of SNr neurons that were inhibited during licking, as well as those with sustained activity during licking. These findings reveal the differential functions of direct and indirect pathway striatal neurons at fine as well as coarse timescales, and support the notion that the two pathways have opponent roles in regulating the basal ganglia output responses during movement.

Orofacial behaviors, such as sniffing, whisking, chewing and licking, are important for sensory exploration and nutrient ingestion (Moore et al., 2014; Travers et al., 1997). Neurons in the striatum are topographically organized (Hintiryan et al., 2016; Hunnicutt et al., 2016; Liu et al., 2020), and the projections from sensorimotor cortical areas representing the mouth region terminate in VLS (Hintiryan et al., 2016). We found that optogenetic inactivation of the VLS, but not the DS, disrupted licking by reducing the number of licks in a bout. This is consistent with previous reports that lesion of VLS impaired the movement of tongue (Pisa, 1988; Pisa and Schranz, 1988), and that stimulation of VLS but not dorsomedial striatum modulated licking (Lee et al., 2020). We showed that inactivation of D1-MSNs and D2-MSNs in the VLS suppressed and increased licking, respectively, consistent with a recent work studying the effect of activating direct and indirect pathways in the VLS (Bakhurin et al., 2020).

In contrast to the opposite roles of direct and indirect pathways observed at behavioral level (Kravitz et al., 2010), cell-type-specific recordings showed that D1- and D2-MSNs are both activated during initiation and execution of voluntary movement (Barbera et al., 2016; Cui et al., 2013; Isomura et al., 2013; Jin et al., 2014; London et al., 2018; Meng et al., 2018; Parker et al., 2018). These findings are consistent with the proposal that the activity of direct pathway facilitates the desired movement and the coactivated indirect pathway inhibits the competing movements (Hikosaka et al., 2000; Jin et al., 2014; Mink, 1996).The direct and indirect pathways are also found to exhibit different activity patterns. For instance, in mice performing learned action sequences of lever pressing, more D1-MSNs displayed sequence-related sustained activity, whereas more D2-MSNs showed sequence-related inhibited activity (Jin et al., 2014). Whether the activity of D1- and D2-MSNs was coactive or opponent also depended on the moment in sequence execution (Geddes et al., 2018). In addition, D1- and D2-MSNs differed in response timing at sub-second timescale during whisker sensorimotor task or showed decorrelated activity in specific epochs of naturalistic behaviors (Markowitz et al., 2018; Sippy et al., 2015). Ex vivo analysis after behavioral training showed that the relative timing between D1- and D2-MSNs switched during transition from goal-directed to habitual behavior (O’Hare et al., 2016). While the previous studies examined MSNs in the dorsal striatum, we studied MSNs in the ventral striatum. We found that D1- and D2-MSNs in the VLS were both active during licking. However, they showed different activity during the initiation and execution of lick sequence. We did not find difference in the percentage of inhibited neurons between D1- and D2-MSNs as reported by a previous study (Jin et al., 2014), which may be due to differences in the behavioral paradigm of movement as well as in the region of striatal recording. Interestingly, for lick-related oscillatory activity on the timescale of a lick cycle (∼ 150 ms), we found significant difference in the phase at peak response between D1- and D2-MSNs. Previous in vivo studies did not observe such difference in relative timing (Cui et al., 2013; Jin et al., 2014; London et al., 2018; Meng et al., 2018; Parker et al., 2018), which may be due to differences in the temporal resolution of response measurements (Ca^2+^ imaging vs. electrophysiology), the temporal characteristics of the movements (lever pressing vs. licking), or the location of striatal regions (dorsal vs. ventral). It should also be noted that, while we used D2-Cre mice, previous studies (Cui et al., 2013; Geddes et al., 2018; London et al., 2018; Meng et al., 2018; Parker et al., 2018) used A2A-Cre mice, the advantage of which over D2-Cre mice is that adenosine A_2A_ receptor does not target cholinergic interneurons in striatum (Durieux et al., 2009).

A previous study found that striatal activity in the direct and indirect pathways was decorrelated at fine timescales, and that the accuracy of decoding behavioral syllable identity was improved when using activity in both pathways as compared to that using activity in each pathway, suggesting that decorrelation between the two pathways is important for representing individual behavioral syllables (Markowitz et al., 2018). In our study, we directly tested the role of fine timescale activity of D1- and D2-MSNs in licking behavior. Intriguingly, we found that transient inactivation of D1-MSNs in VLS around the time of tongue protrusion reduced the spout-contact probability. This demonstrates that the temporally precise activity of D1-MSNs in VLS around tongue protrusion plays a crucial role in controlling individual spout-contact. As the transient impairment of a spout-contact did not affect the probability of next contact, our result also suggests that the innate and stereotyped licking sequence is organized in a hierarchical structure, similar to that found for learned action sequence (Geddes et al., 2018). Furthermore, while inactivation of D2-MSNs for 2 s caused an increase in lick rate, transient inactivation of D2-MSNs around the onset of the third spout-contact or after 3 s following bout onset did not affect licking. This suggests that, compared with the suppression of D1-MSNs activity, it may require a longer duration to inactivate D2-MSNs to drive behavioral effect.

Previous studies showed that activating the GABAergic projection from lateral SNr to superior colliculus suppressed self-initiated licking (Rossi et al., 2016; Toda et al., 2017). Consistent with these reports, we found that inactivation of VLS, which topographically projects to lateral SNr (Lee et al., 2020; von Krosigk et al., 1992), disrupted licking. The GABAergic SNr neurons provide tonic inhibition to downstream areas at the brainstem and thalamus (Hikosaka, 2007). The classical model proposed that the direct and indirect pathways facilitate and decrease movement, respectively, by inhibiting and disinhibiting the responses of SNr neurons (DeLong, 1990; Kravitz et al., 2012). Nevertheless, optogenetic activation of striatal neurons in the direct or indirect pathway caused both decrease and increase in firing rates of SNr neurons (Freeze et al., 2013). Optogenetic inactivation of striatal neurons in the direct or indirect pathway showed that only a small fraction of SNr neurons changed activity, in which both increase and decrease was observed (Tecuapetla et al., 2014). In our study, we found that some SNr neurons displayed inhibited activity and others showed sustained activity during licking, consistent with the proposal that basal ganglia output neurons supporting the desired motor program decrease activity during movement and those involved in competing motor programs increase activity (Hikosaka et al., 2000; Mink, 1996). Interestingly, inactivation of D1-MSNs led to rate increase for a subset of SNr neurons, most of which were inhibited during licking, and rate decrease for another subset, the majority of which showed sustained activity during licking. Inactivation of D2-MSNs also resulted in rate increase and rate decrease for two subsets of SNr neurons, but whose response types were predominantly sustained neurons and inhibited neurons, respectively. The paradoxical observation that inactivation of D1-MSNs (or D2-MSNs) could decrease (or increase) the firing rates of sustained SNr neurons may be due to several possibilities, including the axon collaterals of D1-MSNs in GPe (Gerfen et al., 1990; Xiao et al., 2020), the recurrent collateral connections between D2- and D1-MSNs in the striatum (Taverna et al., 2008), and that sustained neurons (or start neurons) in SNr could inhibit other sustained neurons via axon collaterals within SNr (Brown et al., 2014; Deniau et al., 1982; Deniau et al., 2007). Assuming that the inhibited SNr neurons disinhibit the desired movement, whereas the sustained SNr neurons suppress competing movements, our data suggest that the direct and indirect pathway striatal neurons oppositely control licking movement by differential regulation of SNr activity.

## ACKNOWLEDGMENTS

This work was supported by the Strategic Priority Research Program of Chinese Academy of Sciences (grant No. XDB32010200), Shanghai Municipal Science and Technology Major Project (grant No. 2018SHZDZX05) and the National Natural Science Foundation of China (31771151).

## AUTHOR CONTRIBUTIONS

Z.C., Z-Y. Z. and H. Y. conceived and designed the project. Z.C. and Z-Y. Z. performed most of the experiments and data analysis. T. X. helped with behavioral experiments. W. Z. and X-H. X. performed slice recordings. Y. L. helped with virus injection experiments. H. Y. wrote the manuscript, with feedback from all authors.

## STAR*METHODS

### RESOURCE AVAILABILITY

#### Lead contact

Further information and requests for resources and reagents should be directed to and will be fulfilled by the Lead Contact, Haishan Yao.

## Materials availability

This study did not generate new unique reagents.

## Data and code availability

All data, custom scripts and functions used for this study are available from the Lead Contact, Haishan Yao.

## EXPERIMENTAL MODEL AND SUBJECT DETAILS

### Animals

All animal procedures were approved by the Animal Care and Use Committee at the Institute of Neuroscience, Center for Excellence in Brain Science and Intelligence Technology, Chinese Academy of Sciences (IACUC No. NA-013-2019). We used the following mice: D1-Cre (Tg(Drd1a-cre)262Gsat/Mmcd), D2-Cre (Tg(Drd2-cre)ER44Gsat/Mmcd) (Gong et al., 2007) and C57BL/6 mice. Adult (2–4 months at the time of surgery) male mice were used for all experiments. All mice were housed on a 12 h:12 h light/dark cycle in the Institute of Neuroscience animal facility, with the humidity controlled at 40–70% and temperature at 22–23°C.

## METHOD DETAILS

### Adeno-associated virus

We used the following AAVs: AAV2/5-hSyn-hGtACR1-EGFP-WPRE (titer: 2−3×10^12^ viral particles/ml; for inactivation of VLS or DS); AAV2/8-CAG-DIO-GtACR1-P2A-EGFP (titer: 2−3×10^12^ viral particles/ml; for inactivation of D1-MSNs or D2-MSNs in VLS); AAV2/8-Syn-FLEX-ChrimsonR-tdTomato (titer: 2−3×10^12^ viral particles/ml; for optogenetic tagging of D1- or D2-MSNs); AAV2/8-hSyn-eGFP-3Flag-WPRE-SV40pA or AAV2/8-hSyn-FLEX-tdTomato (titer: 2−3×10^12^ viral particles/ml; for optogenetic experiments as a control group).

### Surgery

Before surgery, the mice were anesthetized with a mixture of fentanyl (0.05 mg/kg), medetomidine (0.5 mg/kg) and midazolam (5 mg/kg) injected intraperitoneally, and were head-fixed in a stereotaxic apparatus. To target VLS or DS, two craniotomies (∼0.5 mm diameter) were performed bilaterally above VLS (AP 0.34 mm, ML ±2.75 mm) or DS (AP 1.0 mm, ML ±1.8 mm). The virus was injected with a glass pipette (15–20 μm tip diameter) using a syringe pump (Harvard Apparatus). We injected 400–500 nl of AAV in VLS at a depth of 3.2 mm and in DS at a depth of 2.0 mm. After the virus injection, the pipette was left in place for 15−30 minutes before retraction. For mice used in optogenetic inactivation experiments, optical fiber (200 μm diameter, NA 0.37) was inserted 200−300 μm above the injection site following the virus injection. A stainless-steel headplate was fixed to the skull using dental cement. For mice used in extracellular recording from VLS or SNr, the skull region above the recording site was marked with permanent ink. After the surgery, the mice were given Rimadyl via drinking water for 3 days, and were allowed to recover with food and water ad libitum for at least 10 days.

In some C57BL/6 mice (n = 6), fluorescently conjugated Cholera toxin subunit B (CTB-555, 2 μg/μl, 300 nl, Invitrogen) was injected unilaterally into VLS. The histology experiments were performed 2 weeks after the injection.

### Behavioral task

Mice were deprived of water for 2 days before the behavioral training. During behavioral experiments, the mouse was head-fixed and sat in an acrylic tube. The lick spout was located around 5 mm in front of the tip of the nose and 0.5 mm below the lower lip. Touch of the spout by forelimbs was prevented by a block of plastic plate. Tongue licks were detected by an electrical lick sensor (Weijnen, 1989) or the interruption of an infrared beam if the mice were used for electrophysiological recordings. Fluid delivery was controlled by a peristaltic valve (Kamoer). An Arduino microcontroller platform was used for spout contact measurement, fluid delivery, and laser stimulation. A multifunction I/O device (USB-6001, National Instruments) was used for data acquisition. The spout contact signals and task-related signals were sampled at 2000 Hz.

For voluntary licking task, mice went through a habituation phase, a free-drinking phase, and a final task phase. During the habituation phase (2–3 days), the mouse was handled by the experimenter for 5–10 min and learned to lick water (300−500 nl) from a syringe. During the free-drinking phase (1–2 days), the mouse was head-fixed into the behavioral apparatus. Water (0.3–0.5 μl) was delivered every time a spout contact was detected. The mouse was head-fixed for at least 30 min, and the lick spout was removed after the mouse consumed 1000 μl of water. During the final task phase, sucrose (10% wt/vol) was used. The delivery of sucrose was triggered if a spout contact was preceded by at least 1 s of no contacts (or 1.5 s of no contacts for electrophysiological recordings of D1- and D2-MSNs), and the time window of spout-contact-triggered delivery of sucrose lasted for 3 s for most experiments and lasted for 4 s for some experiments. For optogenetic inactivation experiments with 2-s laser stimulation or for electrophysiological recording experiments, during the time window of sucrose delivery, 1 μl was delivered after the first spout contact and 0.3–0.5 μl was delivered after each of the following contacts. For optogenetic inactivation experiments with 25-ms or 150-ms laser stimulation, 0.5 μl of sucrose was delivered every 3 contacts. After the time-window of sucrose delivery, there was a waiting period, in which a spout contact would not trigger the delivery of sucrose. The waiting period lasted for 2 s for the experiments of inactivating striatum (VLS or DS) and 1 s for those electrophysiological experiments in which D1- or D2-MSNs were recorded. Each mouse performed the task for 0.5 h in each session. After being trained in the final task phase for 2–3 days, the mice were used for optogenetic or electrophysiological experiments.

### Optogenetic stimulation

For in vivo experiments, optical activation of GtACR1 (ChrimsonR) was induced by green (red) light. A green laser (520 or 532 nm) or a red laser (635 nm) (Shanghai Laser & Optics Century Co.) was connected to an output optical fiber and the laser stimulation was controlled by an Arduino microcontroller. Trials with and without laser stimulation were interleaved.

For optogenetic experiments with 2-s laser stimulation, laser was triggered by the onset of first spout-contact of a bout that was preceded by at least 1 s of no contacts. Laser was at a power of 8 − 10 mW at fiber tip except for that in Figure S6E, which was at a power of 40 mW at fiber tip.

For optogenetic experiments with 25-ms laser stimulation, the inter-contact interval of the mouse was first estimated from a few lick bouts, and laser was triggered after an inter-contact interval following the onset of the second contact in a bout. For optogenetic experiments with 150-ms laser stimulation, laser was triggered at 3 s following the bout onset. For optogenetic inactivation with 25-ms or 150-ms laser stimulation, laser was at a power of 40 mW at fiber tip.

### Optogenetic tagging

For optogenetic tagging of D1- and D2-MSNs, red laser light (1–3 mW at fiber tip) was applied to the fiber of the optrode. We delivered 100 trials of 100 ms laser stimulation, with a 5 s inter-trial interval (Lee et al., 2019). To identify a unit as ChrimsonR-expressing, we required that the unit was significantly activated by laser stimulation, the latency of laser-evoked spikes was < 6 ms and the Pearson’s correlation coefficient between waveforms (1.6 ms duration) of laser-evoked spikes and spontaneous spikes was > 0.95 (Lee et al., 2019). We used a paired t test to compare the spike number between the spikes in a 1-s period before laser onset and the spikes in a 6-ms period after laser onset, and those units with p < 0.01 were considered to be significantly activated by laser stimulation (Lee et al., 2019). These criteria yielded 111 identified ChrimsonR-expressing units in D1-Cre mice (out of 712 total units from 7 mice), 96 identified ChrimsonR-expressing units in D2-Cre mice (out of 784 total units from 6 mice) and 0 false positives (out of 275 total units from 2 control mice) (Figure S4B).

### *In vivo* extracellular recording

Optogenetic tagging and recordings were performed at least 3 weeks after the virus injection. Before the recording, the mice were head-fixed to a holder attached to the stereotaxic apparatus and anesthetized with isoflurane (1–2%). A craniotomy (∼ 1 mm diameter) was made above VLS (AP 0.34 mm, ML ±2.75 mm) or SNr (AP −3.3 to −3.8 mm; ML ±1.65 mm). The dura was removed, and the craniotomy was protected by a silicone elastomer (Kwik-Cast, WPI). The mouse was allowed to recover from the anesthesia in home cage for at least 2 hours. The recordings were made with optrodes (ASSY-37-32-1-9mm, Diagnostic Biochips) or multi-site silicon probes (A1×32-Poly2-10mm-50s-177-A32, NeuroNexus Technologies) mounted on a manipulator (MX7630/45DR, Siskiyou Corporation). The electrodes were coated with DiI or DiO (Invitrogen) to allow post hoc recovery of penetration track. After finishing the recordings, the electrode was retracted. The craniotomy was cleaned with saline and covered with a silicone elastomer (Kwik-Cast, WPI). We made 1–4 sessions of recordings from each mouse.

The neural responses were amplified and filtered using a Cerebus 32-channel system (Blackrock microsystems). Spiking signals were sampled at 30 kHz. To detect the waveforms of spikes, we band-pass filtered the signals at 250–7500 Hz and set a threshold at 3.5 or 4 SD of the background noise. Spikes were sorted offline using the Offline Sorter (Plexon Inc.) based on cluster analysis of principle component amplitude. Spike clusters were considered to be single units if the interspike interval was larger than 1.5 ms and the p value for multivariate analysis of variance tests on clusters was less than 0.05. Task-related events were digitized as TTL levels and recorded by the Cerebus system.

### Slice preparation and recording

We used D1-Cre and D2-Cre mice for slice recordings. Mice that had been injected with AAV2/8-CAG-DIO-GtACR1-P2A-EGFP in the VLS were anesthetized with isoflurane and perfused with ice-cold oxygenated (95% O_2_/5% CO_2_) solution containing the following (in mM): 2.5 KCl, 1.25 NaH_2_PO_4_, 2 Na_2_HPO_4_, 2 MgSO_4_, 213 sucrose, 26 NaHCO_3_. The mouse brain was dissected, and coronal slices including VLS (250 μm) were prepared using a vibratome (VT1200S, Leica) in the ice-cold oxygenated cutting solution. The prepared brain slices were incubated in artificial cerebral spinal fluid (ACSF), which contained the following (in mM): 126 NaCl, 2.5 KCl, 1.25 NaH_2_PO_4_, 1.25 Na_2_HPO_4_, 2 MgSO_4_, 10 Glucose, 26 NaHCO_3_, 2 CaCl_2_, for at least 60 min at 34°C, and then were recorded at ∼33°C with a temperature controller (Warner, TC324B).

D1- or D2-MSNs were identified as EGFP-expressing neurons under fluorescent microscope and visualized by infrared differential interference contrast (BX51, Olympus). Whole-cell recordings in current-clamp mode were made with a Multiclamp 700B amplifier and a Digidata 1440A (Molecular Devices). The electrodes (3−5 MΩ) were filled with an internal solution containing the following (in mM): 135 K-gluconate, 4 KCl, 10 HEPES, 10 sodium phosphocreatine, 4 Mg-ATP, 0.3 Na_3_-GTP and 0.5 biocytin; pH:7.2, 276 mOsm. The spikes of GtACR1-expressiong neurons were induced by holding the membrane potential at −50 ∼ −40 mV with current injection. For optogenetic stimulation, blue or green light (2 s or 25 ms duration, 1−6 mW) was delivered through a ×40 objective using an X-Cite LED light source (Lumen Dynamics). Data were low-pass filtered at 10 kHz and sampled at 10 kHz.

### Histology

The mouse was deeply anesthetized with isoflurane and was perfused with 25 ml saline followed by 25 ml paraformaldehyde (PFA, 4%). Brains were removed, fixed in 4% PFA (4℃) overnight, and then transferred to 30% sucrose in phosphate-buffered saline (PBS) until equilibration. Brains were sectioned at 50 μm using a cryostat (Microm). To observe EGFP fluorescence for brain slices of D1-Cre or D2-Cre mice injected with AAV2/8-CAG-DIO-GtACR1-P2A-EGFP, slices were incubated with blocking buffer (20% BSA, 0.5% Triton X-100 in PBS) for 2 h at room temperature and then with primary antibody (rabbit anti-GFP, 1:1000, Invitrogen, G10362) diluted in blocking buffer overnight at 4℃. Slices were rinsed with PBS and incubated with secondary antibody (donkey anti-rabbit Alexa Fluor 594, 1:1000, Invitrogen, A21207) diluted in blocking buffer for 2 h at room temperature. The sections were rinsed with PBS, mounted onto glass slides and coverslipped with DAPI Fluoromount-G (SouthernBiotech, 0100-20) or Vectashield (Vector, H-1000). Fluorescence images were taken with VS120 (Olympus). Images were analyzed with ImageJ (NIH, US). The atlas schematics in the figures of this study are modified from (Franklin and Paxinos, 2007).

## QUANTIFICATION AND STATISTICAL ANALYSIS

### Data analysis

Analyses were performed in MATLAB. For the licking behavior, we generated timestamps for the onset and offset of each spout contact (0.5 ms resolution) according to the analog voltage signal. The spout contact duration was the time from the onset to the offset of each spout contact (i.e., the time period the tongue contacted the spout). The inter-contact interval was the interval between the onsets of two consecutive contacts. A lick bout was defined as a group of licks in which the first spout contact was preceded by at least 1 s of no contacts, the inter-contact interval of the first three contacts was ≥ 0.3 s, and the last contact was followed by ≥ 0.5 s of no contacts. For voluntary licking task, we constructed peri-stimulus time histogram (PSTH) for the licking behavior by aligning the first contact of all bouts and averaging the licks (200 ms/bin) across bouts. The lick rate in the voluntary licking task was computed using spout contacts within 2 s after bout onset. For the experiments in which 25-ms laser stimulation was applied around the time of tongue protrusion for the third contact, we analyzed spout contact probability for the third and the fourth contact. For the time window of the third (fourth) contact, the onset time was computed as the median time of the third (fourth) contact in laser-off trials minus 50 ms, and the offset time was computed as the median time of the third (fourth) contact in laser-off trials plus 50 ms. Spout contact probability for the third (fourth) contact was defined as the probability that a contact occurred within the time window of the third (fourth) contact.

For the optogenetic tagging experiments, the identified ChrimsonR-expressing units were further classified into putative MSNs, fast-spiking interneurons (FSIs) and tonically active neurons (TANs) based on their spike waveforms, firing rates and coefficients of variation (CV) of inter-spike intervals (Shin et al., 2018). TANs were defined as those with CVs < 1. For non-TAN units, MSNs were defined as those in which peak width > 0.42 ms and peak-valley width > 0.36 ms; FSIs were defined as those in which peak width < 0.42 ms, peak-valley width < 0.36 ms and mean firing rate > 1.5 Hz. Only units classified as putative MSNs and mean firing rate > 0.5 Hz were included in the analysis.

For SNr neurons, we classified the units into GABAergic and non-GABAergic neurons based on their spike waveforms (Figure S7B) (Barter et al., 2015). GABAergic SNr neurons were defined as those with peak width < 0.36 ms. Only units classified as GABAergic SNr neurons and mean firing rate > 1.5 Hz were included in the analysis.

To examine whether the activities of VLS or SNr neurons are related to the start/stop or execution of lick sequence (Jin and Costa, 2010; Jin et al., 2014), we analyzed the spikes for those lick bouts whose length were between 1 s and 4 s. A trial of lick bout was defined to include a baseline phase (from −1 s to −0.5 s relative to bout onset), a start phase (from −0.5 s to 0.25 s relative to bout onset), an execution phase (the duration of the lick bout), and a stop phase (from −0.25 s to 0.5 s relative to bout offset). For both the start and the stop phases, we divided the responses in 30 bins. For the execution phase, as the bout duration varied across different lick bouts during a recording session, we normalized the bout duration by dividing each bout in an equal number of 60 bins (Sales-Carbonell et al., 2018). The binned responses were averaged across all trials to obtain a PSTH. To determine whether the PSTH exhibits significant lick-related activity, we defined an upper (lower) threshold at 3 SD above (below) the mean firing rate in baseline phase. If the firing rates in > 1/3 consecutive bins of the execution phase were above the upper threshold (or below the lower threshold), the neuron was classified as a sustained (or inhibited) cell. For those cells whose responses in the execution phase were not different from the baseline (i.e., neither sustained nor inhibited cells), we defined start (or stop) cells as those whose firing rates in > 1/3 consecutive bins of the start (or stop) phase were above the upper threshold or below the lower threshold, and defined boundary cells as those in which significant change in firing rate was observed for both the start and the stop phases. Those neurons with significant response in at least one of the three phases were considered to exhibit lick-related activity.

To examine whether the VLS neurons shows oscillatory activity related to the lick cycle, we computed spout-contact-triggered spike peri-event time histogram (PETH) using spikes occurring around ±200 ms of each contact, and lick PETH using licks occurring around ±200 ms of each contact within a lick bout (excluding the first two contacts and the last two contacts in the bout) (Rossi et al., 2016). Both the spout-contact-triggered spike PETH and the lick PETH were smoothed with a Gaussian (σ = 25 ms). To determine whether the spout-contact-triggered spike PETH showed oscillatory activity, we computed two Pearson’s correlation coefficients for each unit, one was between the lick PETH and the spout-contact-triggered spike PETH, and the other was between the lick PETH and the spout-contact-triggered spike PETH shifted by 40 ms (corresponding to a phase shift of the spike PETH by ∼ 90°) (10 ms/bin for both lick PETH and spout-contact-triggered spike PETH) (Figure S5). If one of the two coefficients was significant (p < 0.05), the unit was considered to exhibit significant oscillatory activity related to the lick cycle and was used to analyze the oscillatory phase.

To determine the phase relationship between the oscillatory neural activity and the lick cycle, we identified the time at which the amplitude of spout-contact-triggered spike PETH was maximum and transformed it to polar coordinate.

For the responses of SNr neurons with and without inactivation of D1-MSNs (or D2-MSNs) in VLS, we used Wilcoxon signed rank test to compare the firing rates within the first 2 s of lick bout between laser-off and laser-on trials, or compare the firing rates within the time window of the third spout-contact between laser-off and laser-on trials. Those SNr neurons with p < 0.05 were considered to exhibit significant rate increase or rate decrease. Only lick-related SNr neurons were included in this analysis.

## Statistics

The statistical analysis was performed using MATLAB or GraphPad Prism (GraphPad Software). Wilcoxon signed rank test, Wilcoxon rank sum test, χ^2^ test, Rayleigh test, Fisher’s exact test, circular Watson-Williams two-sample test, one-way ANOVA, one-way repeated measures ANOVA, two-way ANOVA with mixed design, or two-way repeated measures ANOVA followed by Sidak’s multiple comparisons test was used to determine the significance of the effect. Correlation values were computed using Pearson’s correlation. Unless otherwise stated, data were reported as SEM.

---

**Figure S1.**
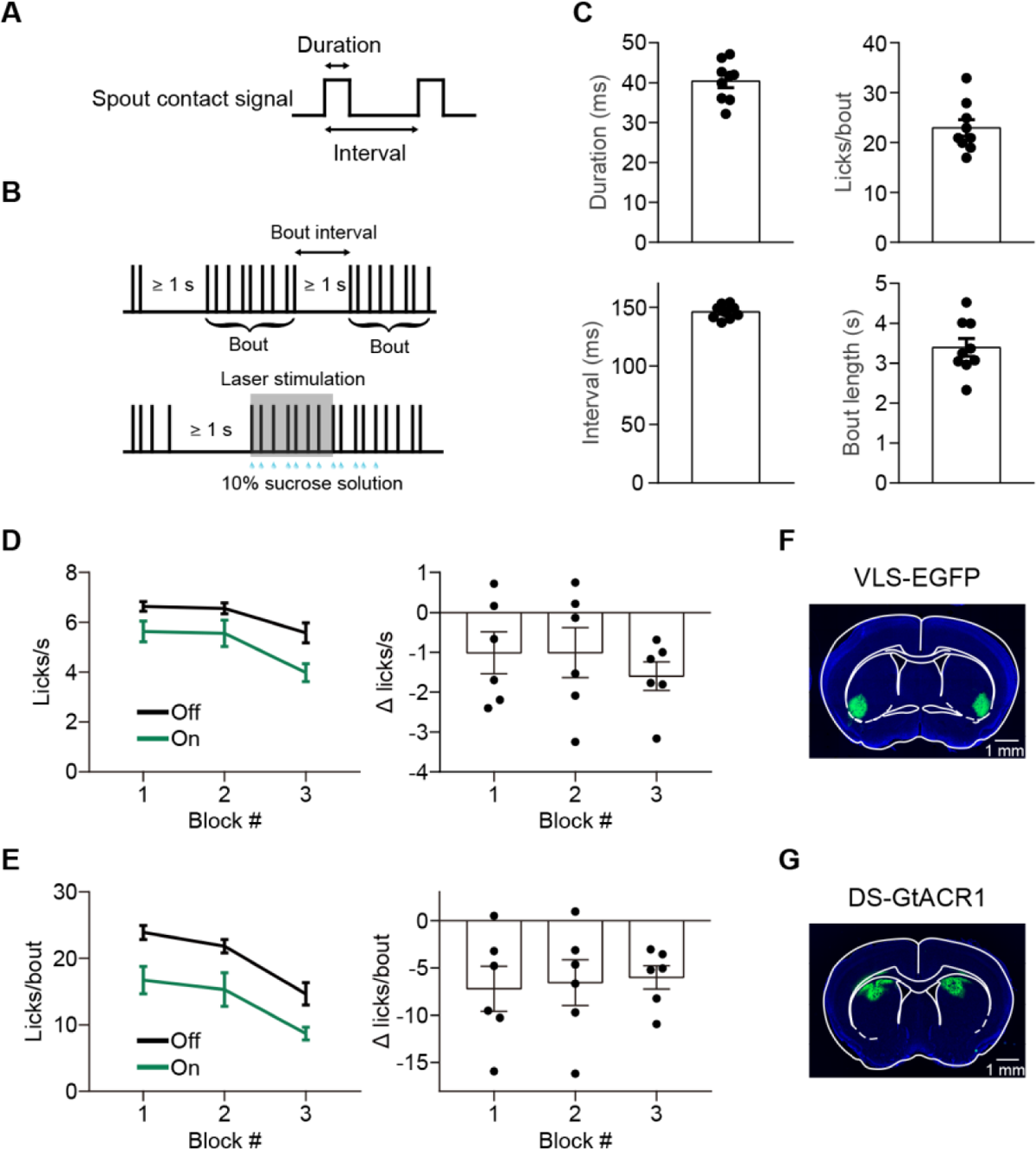
Parameters of Licking Behavior, Effect of Laser Stimulation in Different Blocks of Trials and Fluorescence Images of Virus Injection Sites. Related to Figure 1. (A) Measurement of spout contact duration and inter-contact interval from the spout contact signal. (B) Definition of a valid lick bout and illustration of laser stimulation (gray shading), which was triggered by the first contact of a bout that was preceded by ≥ 1 s of no contacts. (C) Spout contact duration, inter-contact interval, number of licks per bout and bout length for the licking behavior of C57BL/6 mice (n = 9). Data represent mean ± SEM. For these mice, the time window of spout-contact-triggered delivery of sucrose lasted for 4 s. (D) For C57BL/6 mice in which AAV2/5-hSyn-hGtACR1-EGFP-WPRE were bilaterally injected in VLS, trials in the same session were divided into three blocks. Left, lick rates across the three blocks for laser-off and laser-on trials. Right, changes in lick rate (between laser-on and laser-off trials) across the three blocks. F_(1.30, 6.51)_ = 0.6, p = 0.51, one-way repeated measures ANOVA. (E) Analysis for the number of licks per bout, similar to that described in (D). For Δ licks/bout across the three blocks, F_(1.61, 8.04)_ = 0.1, p = 0.86, one-way repeated measures ANOVA. (F) Representative fluorescence image showing the expression of AAV2/8-hSyn-eGFP-3Flag-WPRE-SV40pA in VLS of a C57BL/6 mouse. (G) Representative fluorescence image showing the expression of AAV2/5-hSyn-hGtACR1-EGFP-WPRE in DS of a C57BL/6 mouse.

**Figure S2.**
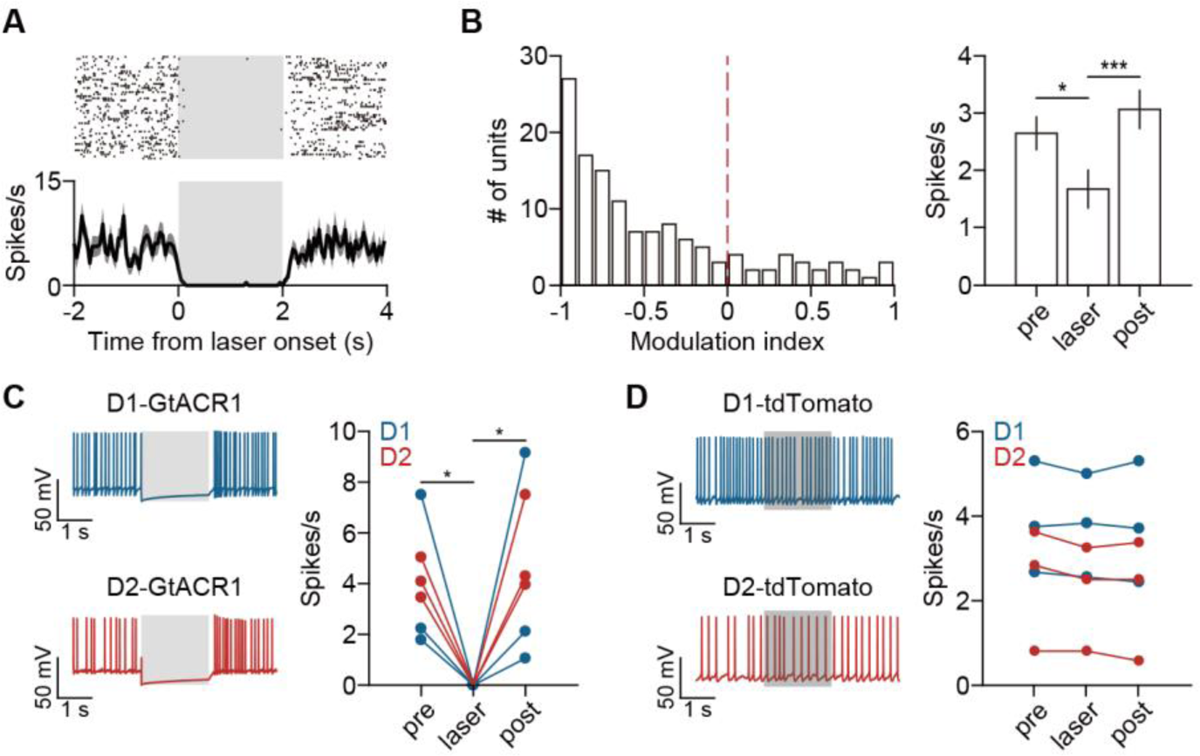
Activation of GtACR1 Suppresses Neuronal Activity in Vivo and in Slices. Related to Figure 1 and Figure 2. (A) Spike raster and PSTH of spontaneous firing for an example VLS neuron recorded from an awake C57BL/6 mouse, in which AAV2/5-hSyn-hGtACR1-EGFP-WPRE was expressed in VLS. Gray shading indicates the duration of laser stimulation. (B) For C57BL/6 mice in which AAV2/5-hSyn-hGtACR1-EGFP-WPRE was expressed in VLS, laser stimulation significantly suppressed the spontaneous firing of VLS neurons. Left, distribution of modulation index, defined as (R_laser_ - R_pre_)/(R_laser_ + R_pre_), in which R_laser_ and R_pre_ are firing rates within a 2-s period during and before laser stimulation, respectively. p = 2.35×10^-13^, Wilcoxon signed rank test. Right, firing rates before, during and after laser stimulation. * p < 0.05, *** p < 0.001, one-way repeated measures ANOVA followed by Sidak’s multiple comparisons test. n = 132 neurons from 2 mice. Data represent mean ± SEM. (C) Left, responses of example D1-MSN and D2-MSN recorded from slices of D1-Cre and D2-Cre mice, in which AAV2/8-CAG-DIO-GtACR1-P2A-EGFP was injected in VLS. Gray shading indicates the duration of laser stimulation. Right, action potentials of D1- and D2-MSNs within a 2-s period before, during and after laser stimulation. * p < 0.05, one-way repeated measures ANOVA followed by Sidak’s multiple comparisons test. n = 6 cells (3 D1-MSNs and 3 D2-MSNs). (D) Left, responses of example D1- and D2-MSNs recorded from slices of D1-Cre and D2-Cre mice, in which AAV2/8-hSyn-FLEX-tdTomato was injected in VLS. Gray shading indicates the duration of laser stimulation. Right, the action potentials of D1- and D2-MSNs, which were induced by current injection, were not significantly affected by laser stimulation. p > 0.05, one-way repeated measures ANOVA. n = 6 cells (3 D1-MSNs and 3 D2-MSNs).

**Figure S3.**
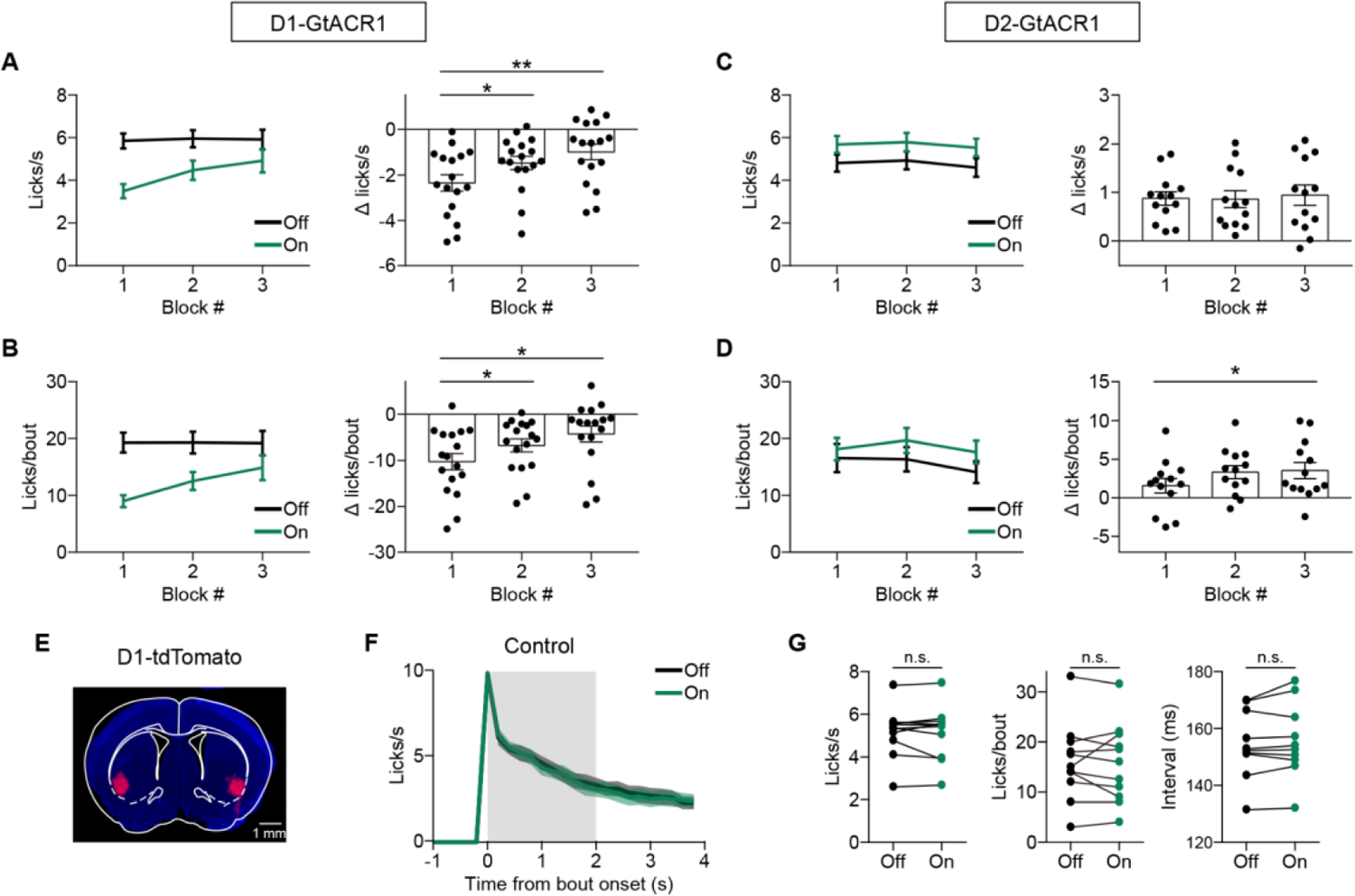
Effect of 2-s Laser Stimulation on Licks in Different Blocks of Trials for D1-Cre or D2-Cre mice and Effect of 2-s Laser Stimulation for Control mice. Related to Figure 2. (A) For D1-Cre mice in which AAV2/8-CAG-DIO-GtACR1-P2A-EGFP were bilaterally injected in VLS, trials in the same session were divided into three blocks. Left, lick rates across the three blocks. Right, changes in lick rate (between laser-on and laser-off trials) across the three blocks. * p < 0.05, ** p < 0.01, one-way repeated measures ANOVA (F_(1.45, 23.18)_ = 9.52, p = 0.0023) followed by Dunnett’s multiple comparisons test. Data represent mean ± SEM. (B) Analysis for the number of licks per bout for D1-Cre mice, similar to that described in (A). For Δ licks/bout across the three blocks, * p < 0.05, one-way repeated measures ANOVA (F_(1.37, 21.89)_ = 8.03, p = 0.0055) followed by Dunnett’s multiple comparisons test. (C) For D2-Cre mice in which AAV2/8-CAG-DIO-GtACR1-P2A-EGFP were bilaterally injected in VLS, trials in the same session were divided into three blocks. Left, lick rates across the three blocks. Right, changes in lick rate (between laser-on and laser-off trials) across the three blocks. F_(1.79, 21.45)_ = 0.16, p = 0.83, one-way repeated measures ANOVA. Data represent mean ± SEM. (D) Analysis for the number of licks per bout for D2-Cre mice, similar to that described in (C). For Δ licks/bout across the three blocks, * p < 0.05, one-way repeated measures ANOVA (F_(1.92, 23.03)_ = 3.88, p = 0.037) followed by Dunnett’s multiple comparisons test. (E) Representative fluorescence image showing the expression of AAV2/8-hSyn-FLEX-tdTomato in VLS of a D1-Cre mouse. (F) Laser stimulation in control D1-Cre mice (n = 5) and control D2-Cre mice (n = 6), in which AAV2/8-hSyn-FLEX-tdTomato was bilaterally injected in VLS, did not change the lick PSTH (F_(1, 10)_ = 0.13, p = 0.72, two-way repeated measures ANOVA). Data represent mean ± SEM. Gray shading indicates the duration of laser stimulation. (G) Laser stimulation in control D1-Cre mice (n = 5) and control D2-Cre mice (n = 6), in which AAV2/8-hSyn-FLEX-tdTomato was bilaterally injected in VLS, did not change lick rate (p = 0.97), number of licks per bout (p = 0.55) or inter-contact interval (p = 0.24, Wilcoxon signed rank test).

**Figure S4.**
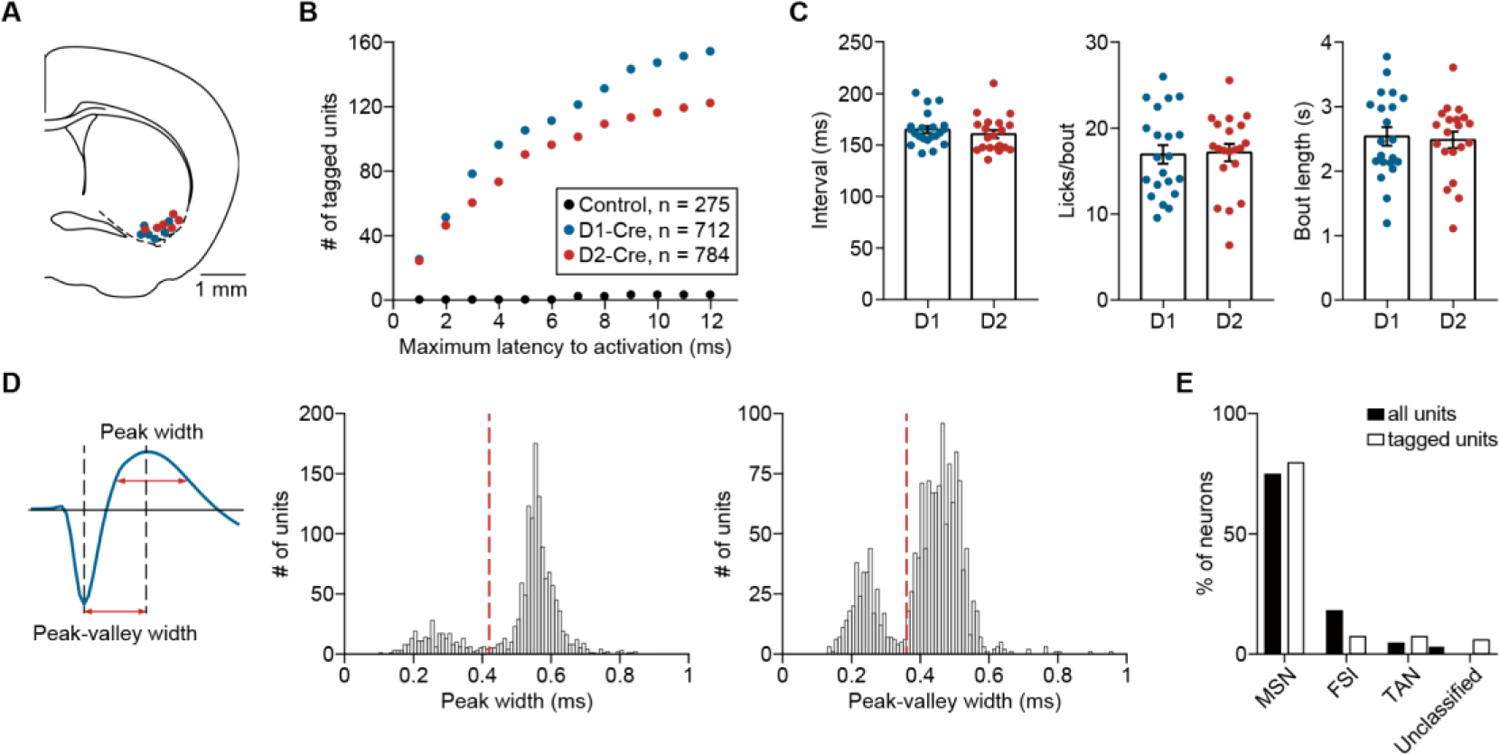
Spike Waveform Parameters of Units Identified by Optogenetic Tagging from the VLS of D1-Cre and D2-Cre Mice. Related to Figure 3. (A) Each dot represents the location of the electrode tip in the last recording session for each mouse. Blue, sites of electrode tip for D1-Cre mice. Red, sites of electrode tip for D2-Cre mice. The distance between the uppermost and the lowermost recording sites in the electrode was 300 μm. (B) Cumulative number of units identified by optogenetic tagging as a function of maximum latency to activation. Total number of cells for D1-Cre, D2-Cre and control (C57BL/6) mice were 712, 784, and 275, respectively. (C) Licking parameters were not significantly different between D1-Cre and D2-Cre mice used in the optogenetic tagging experiments. Inter-contact interval: p = 0.43; number of licks per bout: p = 0.72; bout length: p = 0.84, Wilcoxon rank sum test. D1: n = 21 sessions from 6 D1-Cre mice; D2: n = 20 sessions from 6 D2-Cre mice. Data represent mean ± SEM. (D) Left, illustration of the spike waveform parameters. Middle, Distribution of peak width for all units (n = 1496) recorded in the optogenetic tagging experiments. Right, Distribution of peak-valley width for all recorded units (n = 1496). For non-TAN units, MSNs were defined as those in which peak width > 0.42 ms (vertical dashed line in the middle panel) and peak-valley width > 0.36 ms (vertical dashed line in the right panel). (E) Proportion of MSNs, FSIs, TANs and unclassified units among all recorded units (n = 1496) and tagged units (n = 207).

**Figure S5.**
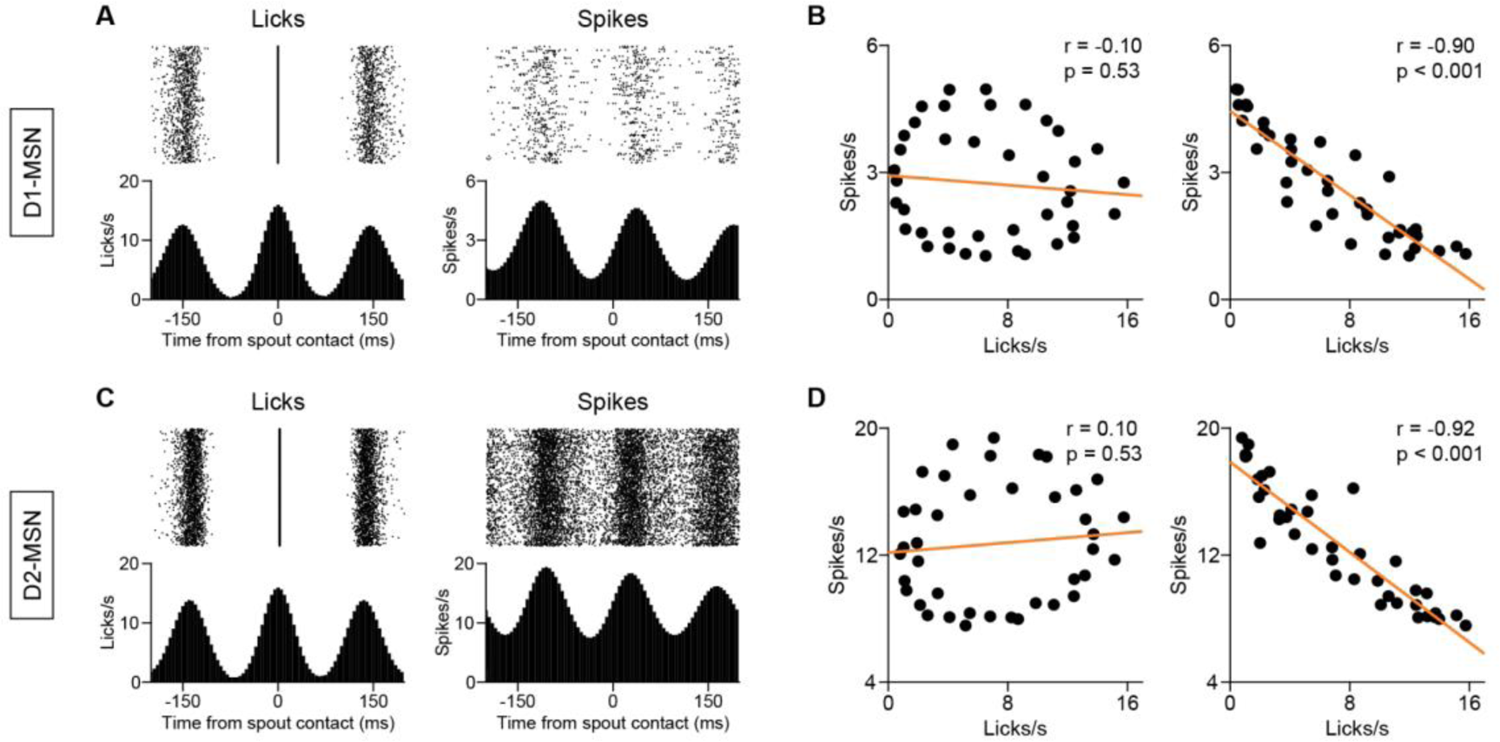
Correlation between Lick PETH and Spout-contact-triggered Spike PETH. Related to Figure 4. (A) Left, lick rasters and lick PETH computed using licks around ±200 ms of each spout-contact, for an example D1-Cre mouse. Right, spout-contact-triggered spikes (rasters and PETH), computed using spikes occurring around ±200 ms of each contact, for an example D1-MSN recorded from the mouse shown in the left. Both the lick PETH and the spout-contact-triggered spike PETH were smoothed with a Gaussian (σ = 25 ms). (B) Left, correlation between lick PETH and spout-contact-triggered spike PETH in (A). Right, correlation between lick PETH in (A) and spout-contact-triggered spike PETH in (A) shifted by 40 ms (corresponding to a phase shift of ∼90°). As one of the two coefficients was significant (p < 0.05), this unit was considered to exhibit significant oscillatory activity related to the lick cycle. (C) Left, lick rasters and lick PETH computed using licks around ±200 ms of each spout-contact, for an example D2-Cre mouse. Right, spout-contact-triggered spikes (rasters and spike PETH), computed using spikes occurring around ±200 ms of each contact, for an example D2-MSN from the mouse shown in the left. Similar to that described in (A). (D) Left, correlation between lick PETH and spout-contact-triggered spike PETH in (C). Right, correlation between lick PETH in (C) and spout-contact-triggered spike PETH in (C) shifted by 40 ms (corresponding to a phase shift of ∼90°). Similar to that described in (B).

**Figure S6.**
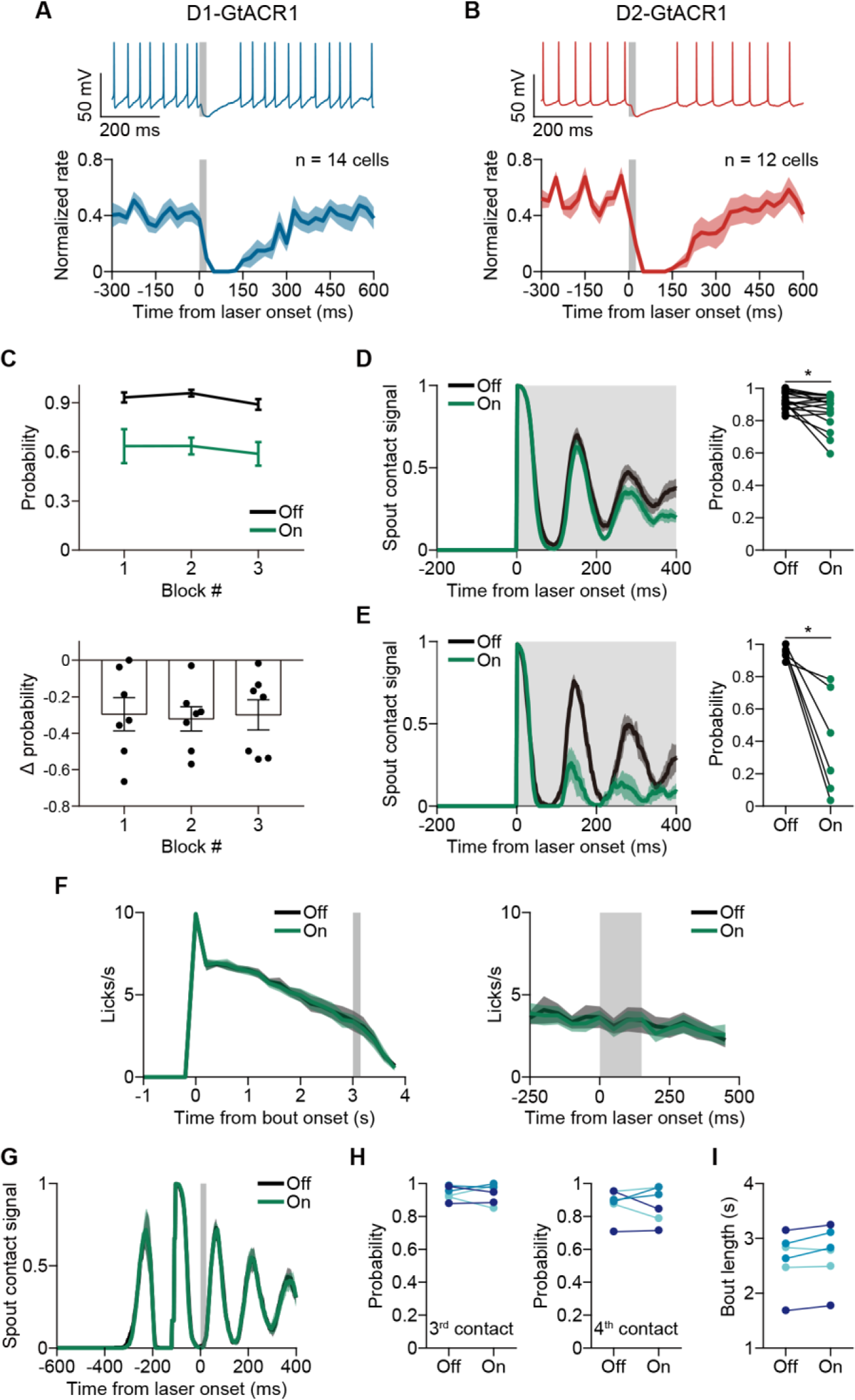
Effect of Transient Laser Stimulation on Spiking of D1- and D2-MSNs, and on Licking Behavior in D1-GtACR1 Mice, D2-GtACR1 Mice and Control Mice. Related to Figure 5. (A) Upper, slice recording of an example D1-MSN from a D1-Cre mouse, in which AAV2/8-CAG-DIO-GtACR1-P2A-EGFP was injected in VLS. Lower, effect of 25-ms laser stimulation on the responses of population D1-MSNs (n = 14 from 4 mice). For each neuron, the responses were normalized by the peak firing rate in [-300 0] ms before laser onset. Data represent mean ± SEM. Gray shading, duration of laser stimulation. (B) Upper, slice recording of an example D2-MSN from a D2-Cre mouse, in which AAV2/8-CAG-DIO-GtACR1-P2A-EGFP was injected in VLS. Lower, effect of 25-ms laser stimulation on the responses of population D2-MSNs (n = 12 from 4 mice). Similar to that described in (A). (C) To analyze the effect of transient laser stimulation on spout contact probability for D1-Cre mice in which AAV2/8-CAG-DIO-GtACR1-P2A-EGFP were bilaterally injected in VLS, trials in the same session were divided into three blocks. Laser stimulation (duration = 25 ms) was triggered after an inter-contact interval following the onset of the second contact. Upper, probability of third spout-contact across the three blocks. Lower, the reduction in the probability of third spout-contact did not differ across the 3 blocks. F_(1.85, 11.11)_ = 0.052, p = 0.94, one-way repeated measures ANOVA. (D) Left, inactivation of D1-MSNs (with 2-s laser stimulation at 8−10 mW) caused a small but significant reduction in the amplitude of spout contact signal for the contact immediately after laser onset (F_(1, 16)_ = 47.5, p = 3.62×10^-6^, two-way repeated measures ANOVA). Gray shading, duration of laser stimulation, which was triggered by the onset of first spout-contact. Data represent mean ± SEM. Right, inactivation of D1-MSNs (with 2-s laser stimulation at 8−10 mW) caused a small but significant reduction in the spout contact probability for the contact immediately after laser onset. * p < 0.05, n = 17 mice, Wilcoxon signed rank test. (E) Left, inactivation of D1-MSNs (2-s laser at 40 mW) significantly reduced the amplitude of spout contact signal for the contact immediately after laser onset (F_(1, 5)_ = 45.09, p = 1.11×10^-3^, two-way repeated measures ANOVA). Gray shading, duration of laser stimulation, which was triggered by the onset of first spout-contact. Data represent mean ± SEM. Right, inactivation of D1-MSNs (2-s laser at 40 mW) significantly reduced the spout contact probability for the contact immediately after laser onset. * p < 0.05, n = 6 mice, Wilcoxon signed rank test. (F) Inactivating D2-MSNs for 150 ms (laser power at 40 mW) at 3 s after bout onset, when lick rate was lower, did not affect licking behavior. F_(1, 6)_ = 0.18, p = 0.69, two-way repeated measures ANOVA. Data represent mean ± SEM. Gray shading indicates the duration of laser stimulation. (G) Transient laser stimulation did not change the amplitude of spout contact signal for control D1-Cre mice in which AAV2/8-hSyn-FLEX-tdTomato was injected. F_(1, 5)_ = 0.69, p = 0.45, two-way repeated measures ANOVA. Data represent mean ± SEM. The spout contact signals were aligned to laser onset. Gray shading indicates the duration of laser stimulation. (H) Transient laser stimulation did not change the spout contact probability for control D1-Cre mice in which AAV2/8-hSyn-FLEX-tdTomato was injected. Left, for the third contact, p = 1, Wilcoxon signed rank test. Right, for the fourth contact, p = 1, Wilcoxon signed rank test. n = 6 sessions from 3 mice (each color represents one mouse). (I) Transient laser stimulation did not change bout length for control D1-Cre mice in which AAV2/8-hSyn-FLEX-tdTomato was injected. p = 0.094, Wilcoxon signed rank test. n = 6 sessions from 3 mice (each color represents one mouse).

**Figure S7.**
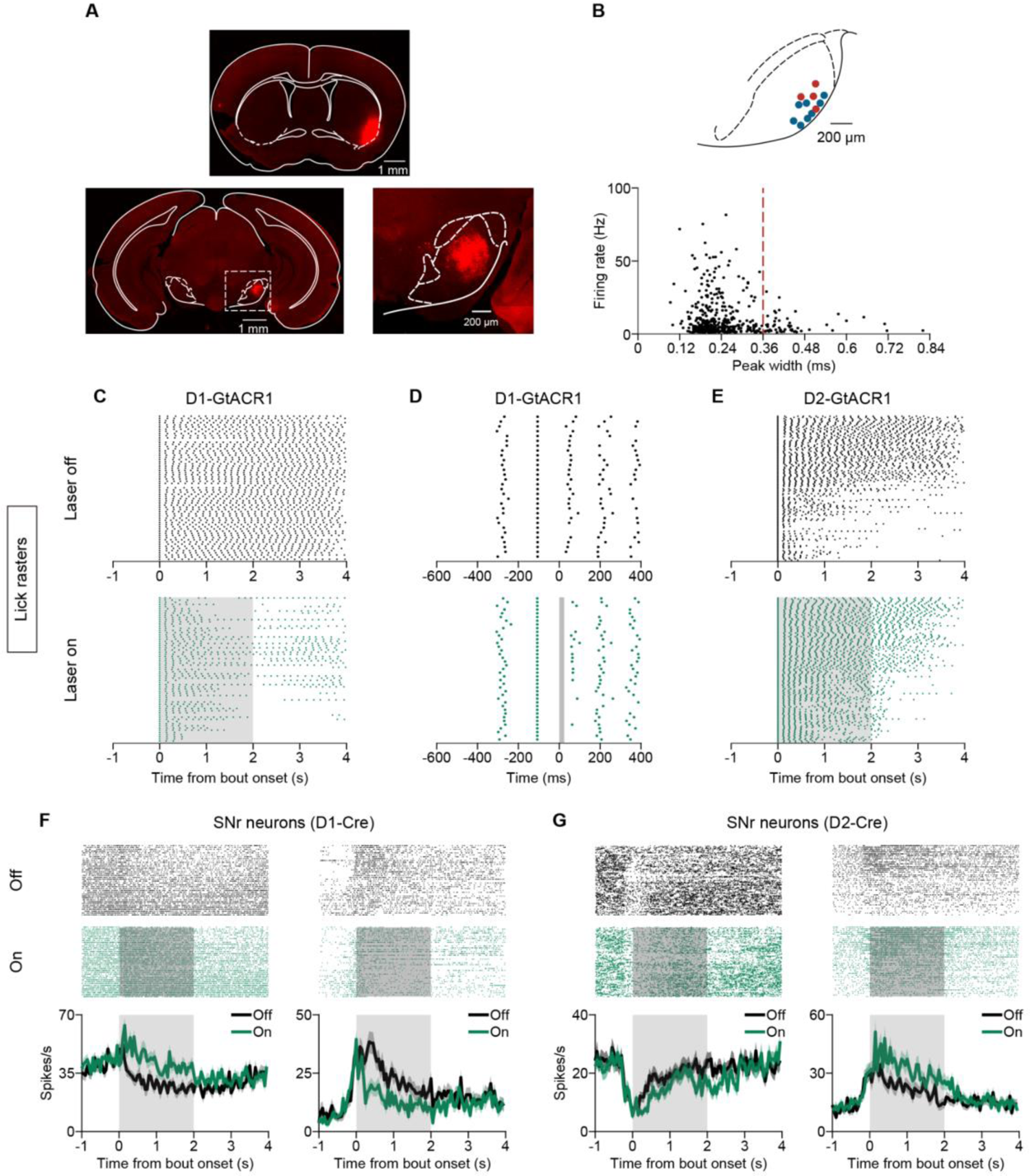
Recording from SNr while Inactivation of D1- or D2-MSNs in VLS. Related to Figure 6 and Figure 7. (A) CTB-555 was injected in VLS (upper) and axon terminals from VLS were found in lateral part of SNr (lower). Lower right, enlarged view of image in the white box. (B) Upper, each dot represents the location of electrode tip in the last recording session for each mouse. Blue, sites of electrode tip for D1-Cre mice. Red, sites of electrode tip for D2-Cre mice. The distance between the uppermost and the lowermost recording sites in the electrode was 775 μm. Lower, firing rate versus peak width of the SNr neurons (n = 512). Units with peak width < 0.36 ms were considered as GABAergic neurons. (C) Lick rasters for laser-off and laser-on trials of an example D1-Cre mouse, in which AAV2/8-CAG-DIO-GtACR1-P2A-EGFP was bilaterally injected in VLS. Gray shading indicates duration of 2-s laser stimulation. (D) Lick rasters for laser-off and laser-on trials of an example D1-Cre mouse, in which AAV2/8-CAG-DIO-GtACR1-P2A-EGFP was bilaterally injected in VLS. After the first spout-contact, 25-ms of laser stimulation (gray shading) was triggered after an inter-contact interval following the onset of the second contact. (E) Lick rasters for laser-off and laser-on trials of an example D2-Cre mouse, in which AAV2/8-CAG-DIO-GtACR1-P2A-EGFP was bilaterally injected in VLS. Gray shading indicates duration of 2-s laser stimulation. (F) Inactivation of D1-MSNs in VLS by 2-s laser stimulation caused an increase in firing rate in an example SNr neuron (left) and a decrease in firing rate in another example SNr neuron (right). Data represent mean ± SEM. Gray shading indicates duration of laser stimulation. (G) Inactivation of D2-MSNs in VLS by 2-s laser stimulation caused a decrease in firing rate in an example SNr neuron (left) and an increase in firing rate in another example SNr neuron (right). Data represent mean ± SEM. Gray shading indicates duration of laser stimulation.

## REFERENCES

1. Albin, R.L., Young, A.B., and Penney, J.B. (1989). The functional anatomy of basal ganglia disorders. Trends Neurosci. 12, 366–375.

2. Aldridge, J.W., and Berridge, K.C. (1998). Coding of serial order by neostriatal neurons: a “natural action” approach to movement sequence. J. Neurosci. 18, 2777–2787.

3. Bakhurin, K.I., Li, X., Friedman, A.D., Lusk, N.A., Watson, G.D., Kim, N., and Yin, H.H. (2020). Opponent regulation of action performance and timing by striatonigral and striatopallidal pathways. Elife 9, e54831.

4. Bakhurin, K.I., Mac, V., Golshani, P., and Masmanidis, S.C. (2016). Temporal correlations among functionally specialized striatal neural ensembles in reward-conditioned mice. J. Neurophysiol. 115, 1521–1532.

5. Barbera, G., Liang, B., Zhang, L., Gerfen, C.R., Culurciello, E., Chen, R., Li, Y., and Lin, D.T. (2016). Spatially Compact Neural Clusters in the Dorsal Striatum Encode Locomotion Relevant Information. Neuron 92, 202–213.

6. Barter, J.W., Li, S., Sukharnikova, T., Rossi, M.A., Bartholomew, R.A., and Yin, H.H. (2015). Basal ganglia outputs map instantaneous position coordinates during behavior. J. Neurosci. 35, 2703–2716.

7. Berens, P. (2009). CircStat: A MATLAB Toolbox for Circular Statistics. J Stat Softw 31, 1–21.

8. Brown, J., Pan, W.X., and Dudman, J.T. (2014). The inhibitory microcircuit of the substantia nigra provides feedback gain control of the basal ganglia output. Elife 3, e02397.

9. Cui, G., Jun, S.B., Jin, X., Pham, M.D., Vogel, S.S., Lovinger, D.M., and Costa, R.M. (2013). Concurrent activation of striatal direct and indirect pathways during action initiation. Nature 494, 238–242.

10. DeLong, M.R. (1990). Primate models of movement disorders of basal ganglia origin. Trends Neurosci. 13, 281–285.

11. Deniau, J.M., Kitai, S.T., Donoghue, J.P., and Grofova, I. (1982). Neuronal interactions in the substantia nigra pars reticulata through axon collaterals of the projection neurons. An electrophysiological and morphological study. Exp. Brain Res. 47, 105–113.

12. Deniau, J.M., Mailly, P., Maurice, N., and Charpier, S. (2007). The pars reticulata of the substantia nigra: a window to basal ganglia output. Prog. Brain Res. 160, 151–172.

13. Durieux, P.F., Bearzatto, B., Guiducci, S., Buch, T., Waisman, A., Zoli, M., Schiffmann, S.N., and de Kerchove d’Exaerde, A. (2009). D2R striatopallidal neurons inhibit both locomotor and drug reward processes. Nat. Neurosci. 12, 393–395.

14. Fan, D., Rossi, M.A., and Yin, H.H. (2012). Mechanisms of action selection and timing in substantia nigra neurons. J. Neurosci. 32, 5534–5548.

15. Franklin, K.B.J., and Paxinos, G. (2007). The mouse brain in stereotaxic coordinates, 3rd Edition (Amsterdam, Boston: Elsevier Academic Press).

16. Freeze, B.S., Kravitz, A.V., Hammack, N., Berke, J.D., and Kreitzer, A.C. (2013). Control of basal ganglia output by direct and indirect pathway projection neurons. J. Neurosci. 33, 18531–18539.

17. Geddes, C.E., Li, H., and Jin, X. (2018). Optogenetic Editing Reveals the Hierarchical Organization of Learned Action Sequences. Cell 174, 32–43 e15.

18. Gerfen, C.R., Engber, T.M., Mahan, L.C., Susel, Z., Chase, T.N., Monsma, F.J., Jr., and Sibley, D.R. (1990). D1 and D2 dopamine receptor-regulated gene expression of striatonigral and striatopallidal neurons. Science 250, 1429–1432.

19. Gong, S., Doughty, M., Harbaugh, C.R., Cummins, A., Hatten, M.E., Heintz, N., and Gerfen, C.R. (2007). Targeting Cre recombinase to specific neuron populations with bacterial artificial chromosome constructs. J. Neurosci. 27, 9817–9823.

20. Gulley, J.M., Kosobud, A.E., and Rebec, G.V. (2002). Behavior-related modulation of substantia nigra pars reticulata neurons in rats performing a conditioned reinforcement task. Neuroscience 111, 337–349.

21. Hikosaka, O. (2007). Basal ganglia mechanisms of reward-oriented eye movement. Ann N Y Acad Sci 1104, 229–249.

22. Hikosaka, O., Takikawa, Y., and Kawagoe, R. (2000). Role of the basal ganglia in the control of purposive saccadic eye movements. Physiol. Rev. 80, 953–978.

23. Hintiryan, H., Foster, N.N., Bowman, I., Bay, M., Song, M.Y., Gou, L., Yamashita, S., Bienkowski, M.S., Zingg, B., Zhu, M., et al. (2016). The mouse cortico-striatal projectome. Nat. Neurosci. 19, 1100–1114.

24. Hunnicutt, B.J., Jongbloets, B.C., Birdsong, W.T., Gertz, K.J., Zhong, H., and Mao, T. (2016). A comprehensive excitatory input map of the striatum reveals novel functional organization. Elife 5, e19103.

25. Isomura, Y., Takekawa, T., Harukuni, R., Handa, T., Aizawa, H., Takada, M., and Fukai, T. (2013). Reward-modulated motor information in identified striatum neurons. J. Neurosci. 33, 10209–10220.

26. Jin, X., and Costa, R.M. (2010). Start/stop signals emerge in nigrostriatal circuits during sequence learning. Nature 466, 457–462.

27. Jin, X., and Costa, R.M. (2015). Shaping action sequences in basal ganglia circuits. Curr. Opin. Neurobiol. 33, 188–196.

28. Jin, X., Tecuapetla, F., and Costa, R.M. (2014). Basal ganglia subcircuits distinctively encode the parsing and concatenation of action sequences. Nat. Neurosci. 17, 423–430.

29. Klaus, A., Alves da Silva, J., and Costa, R.M. (2019). What, If, and When to Move: Basal Ganglia Circuits and Self-Paced Action Initiation. Annu. Rev. Neurosci. 42, 459–483.

30. Kravitz, A.V., Freeze, B.S., Parker, P.R., Kay, K., Thwin, M.T., Deisseroth, K., and Kreitzer, A.C. (2010). Regulation of parkinsonian motor behaviours by optogenetic control of basal ganglia circuitry. Nature 466, 622–626.

31. Kravitz, A.V., Tye, L.D., and Kreitzer, A.C. (2012). Distinct roles for direct and indirect pathway striatal neurons in reinforcement. Nat. Neurosci. 15, 816–818.

32. Lee, J., Wang, W., and Sabatini, B.L. (2020). Anatomically segregated basal ganglia pathways allow parallel behavioral modulation. Nat. Neurosci. 23, 1388–1398.

33. Lee, K., Bakhurin, K.I., Claar, L.D., Holley, S.M., Chong, N.C., Cepeda, C., Levine, M.S., and Masmanidis, S.C. (2019). Gain Modulation by Corticostriatal and Thalamostriatal Input Signals during Reward-Conditioned Behavior. Cell Rep 29, 2438–2449 e2434.

34. Liu, D., Li, W., Ma, C., Zheng, W., Yao, Y., Tso, C.F., Zhong, P., Chen, X., Song, J.H., Choi, W., et al. (2020). A common hub for sleep and motor control in the substantia nigra. Science 367, 440–445.

35. London, T.D., Licholai, J.A., Szczot, I., Ali, M.A., LeBlanc, K.H., Fobbs, W.C., and Kravitz, A.V. (2018). Coordinated Ramping of Dorsal Striatal Pathways preceding Food Approach and Consumption. J. Neurosci. 38, 3547–3558.

36. Markowitz, J.E., Gillis, W.F., Beron, C.C., Neufeld, S.Q., Robertson, K., Bhagat, N.D., Peterson, R.E., Peterson, E., Hyun, M., Linderman, S.W., et al. (2018). The Striatum Organizes 3D Behavior via Moment-to-Moment Action Selection. Cell 174, 44–58 e17.

37. Martiros, N., Burgess, A.A., and Graybiel, A.M. (2018). Inversely Active Striatal Projection Neurons and Interneurons Selectively Delimit Useful Behavioral Sequences. Curr. Biol. 28, 560–573 e565.

38. Meng, C., Zhou, J., Papaneri, A., Peddada, T., Xu, K., and Cui, G. (2018). Spectrally Resolved Fiber Photometry for Multi-component Analysis of Brain Circuits. Neuron 98, 707–717 e704.

39. Mink, J.W. (1996). The basal ganglia: focused selection and inhibition of competing motor programs. Prog. Neurobiol. 50, 381–425.

40. Mittler, T., Cho, J., Peoples, L.L., and West, M.O. (1994). Representation of the body in the lateral striatum of the freely moving rat: single neurons related to licking. Exp. Brain Res. 98, 163–167.

41. Moore, J.D., Kleinfeld, D., and Wang, F. (2014). How the brainstem controls orofacial behaviors comprised of rhythmic actions. Trends Neurosci. 37, 370–380.

42. Nelson, A.B., and Kreitzer, A.C. (2014). Reassessing models of basal ganglia function and dysfunction. Annu. Rev. Neurosci. 37, 117–135.

43. O’Hare, J.K., Ade, K.K., Sukharnikova, T., Van Hooser, S.D., Palmeri, M.L., Yin, H.H., and Calakos, N. (2016). Pathway-Specific Striatal Substrates for Habitual Behavior. Neuron 89, 472–479.

44. Oldenburg, I.A., and Sabatini, B.L. (2015). Antagonistic but Not Symmetric Regulation of Primary Motor Cortex by Basal Ganglia Direct and Indirect Pathways. Neuron 86, 1174–1181.

45. Park, J., Coddington, L.T., and Dudman, J.T. (2020). Basal Ganglia Circuits for Action Specification. Annu. Rev. Neurosci. 43, 485–507.

46. Parker, J.G., Marshall, J.D., Ahanonu, B., Wu, Y.W., Kim, T.H., Grewe, B.F., Zhang, Y., Li, J.Z., Ding, J.B., Ehlers, M.D., and Schnitzer, M.J. (2018). Diametric neural ensemble dynamics in parkinsonian and dyskinetic states. Nature 557, 177–182.

47. Pisa, M. (1988). Motor functions of the striatum in the rat: critical role of the lateral region in tongue and forelimb reaching. Neuroscience 24, 453–463.

48. Pisa, M., and Schranz, J.A. (1988). Dissociable motor roles of the rat’s striatum conform to a somatotopic model. Behav. Neurosci. 102, 429–440.

49. Redgrave, P., Prescott, T.J., and Gurney, K. (1999). The basal ganglia: a vertebrate solution to the selection problem? Neuroscience 89, 1009–1023.

50. Rossi, M.A., Li, H.E., Lu, D., Kim, I.H., Bartholomew, R.A., Gaidis, E., Barter, J.W., Kim, N., Cai, M.T., Soderling, S.H., and Yin, H.H. (2016). A GABAergic nigrotectal pathway for coordination of drinking behavior. Nat. Neurosci. 19, 742–748.

51. Rossi, M.A., and Yin, H.H. (2015). Elevated dopamine alters consummatory pattern generation and increases behavioral variability during learning. Front Integr Neurosci 9, 37.

52. Sales-Carbonell, C., Taouali, W., Khalki, L., Pasquet, M.O., Petit, L.F., Moreau, T., Rueda-Orozco, P.E., and Robbe, D. (2018). No Discrete Start/Stop Signals in the Dorsal Striatum of Mice Performing a Learned Action. Curr. Biol. 28, 3044–3055 e3045.

53. Shin, J.H., Kim, D., and Jung, M.W. (2018). Differential coding of reward and movement information in the dorsomedial striatal direct and indirect pathways. Nat. Commun. 9, 404.

54. Sippy, T., Lapray, D., Crochet, S., and Petersen, C.C. (2015). Cell-Type-Specific Sensorimotor Processing in Striatal Projection Neurons during Goal-Directed Behavior. Neuron 88, 298–305.

55. Smith, Y., Bevan, M.D., Shink, E., and Bolam, J.P. (1998). Microcircuitry of the direct and indirect pathways of the basal ganglia. Neuroscience 86, 353–387.

56. Taverna, S., Ilijic, E., and Surmeier, D.J. (2008). Recurrent collateral connections of striatal medium spiny neurons are disrupted in models of Parkinson’s disease. J. Neurosci. 28, 5504–5512.

57. Tecuapetla, F., Jin, X., Lima, S.Q., and Costa, R.M. (2016). Complementary Contributions of Striatal Projection Pathways to Action Initiation and Execution. Cell 166, 703–715.

58. Tecuapetla, F., Matias, S., Dugue, G.P., Mainen, Z.F., and Costa, R.M. (2014). Balanced activity in basal ganglia projection pathways is critical for contraversive movements. Nat. Commun. 5, 4315.

59. Toda, K., Lusk, N.A., Watson, G.D.R., Kim, N., Lu, D., Li, H.E., Meck, W.H., and Yin, H.H. (2017). Nigrotectal Stimulation Stops Interval Timing in Mice. Curr. Biol. 27, 3763–3770 e3763.

60. Travers, J.B., Dinardo, L.A., and Karimnamazi, H. (1997). Motor and premotor mechanisms of licking. Neurosci Biobehav Rev 21, 631–647.

61. Vandaele, Y., Mahajan, N.R., Ottenheimer, D.J., Richard, J.M., Mysore, S.P., and Janak, P.H. (2019). Distinct recruitment of dorsomedial and dorsolateral striatum erodes with extended training. Elife 8, e49536.

62. von Krosigk, M., Smith, Y., Bolam, J.P., and Smith, A.D. (1992). Synaptic organization of GABAergic inputs from the striatum and the globus pallidus onto neurons in the substantia nigra and retrorubral field which project to the medullary reticular formation. Neuroscience 50, 531–549.

63. Weijnen, J.A. (1989). Lick sensors as tools in behavioral and neuroscience research. Physiol. Behav. 46, 923–928.

64. Weijnen, J.A. (1998). Licking behavior in the rat: measurement and situational control of licking frequency. Neurosci Biobehav Rev 22, 751–760.

65. Xiao, X., Deng, H., Furlan, A., Yang, T., Zhang, X., Hwang, G.R., Tucciarone, J., Wu, P., He, M., Palaniswamy, R., et al. (2020). A Genetically Defined Compartmentalized Striatal Direct Pathway for Negative Reinforcement. Cell 183, 211–227 e220.

66. Yttri, E.A., and Dudman, J.T. (2016). Opponent and bidirectional control of movement velocity in the basal ganglia. Nature 533, 402–406.

